# Impacts of species introductions on the trait diversity of interacting avian frugivores and fleshy-fruited plants depend on native trait diversity

**DOI:** 10.1101/2025.01.18.633718

**Authors:** Larissa Nowak, Evan C. Fricke, Anna Traveset, Isabel Donoso

## Abstract

1. Introductions of non-native species to native ecological communities by humans are major drivers of global biodiversity change. At the same time, biotic interactions, such as those between avian frugivores and fleshy-fruited plants, build the backbone of ecosystems. Hence, studying how species introductions influence interacting species of both trophic levels is essential to understand possible functional consequences of global change.
2. Here, we take a trait-based approach, focusing on species traits that influence their biotic interactions and related ecosystem functioning, and explore how species introductions affect the diversity of such functional traits within and across assemblages of interacting fleshy-fruited plants and frugivorous birds at several locations across the globe.
3. Specifically, we calculate differences in trait diversity and mean trait values, and compare beta trait diversity with and without introduced species, for 25 frugivorous bird and 62 fleshy-fruited plant assemblages.
4. Introduced species tended to increase bird and plant trait diversity in assemblages with low native trait diversity but decreased it in assemblages with higher native trait diversity. In bird assemblages, species introductions shifted mean values towards larger birds with wider bills and more pointed wings. In plant assemblages, mean trait values showed variable changes: fruit length increased and plant height decreased.
5. Comparisons between assemblages revealed that introduced species reduced the beta trait diversity of both frugivorous birds and fleshy-fruited plants, suggesting biotic homogenisation in terms of their functional traits.
6. Changes in trait diversity underscore that species introductions can have functional consequences for biotic interactions and related ecosystem functions, potentially affecting the availability of interaction partners with matching traits and the provisioning of seed dispersal.

## Introduction

Global change rapidly alters biodiversity (Díaz et al., 2019). Studying biodiversity and ecosystem functions under global change requires considering species integration into complex assemblages of interacting species as biotic interactions are the foundation of ecosystems and can mediate and accelerate global change effects on biodiversity (Brodie et al., 2014; Schleuning et al., 2020). Trait-based approaches provide a promising way to study assemblages of interacting species because species traits influence abiotic and biotic interactions and species’ functional roles in ecosystems (Bello et al., 2023; Tilman, 2001).

An important driver of global biodiversity change is the introduction of non-native (or alien) species to native ecological communities (Díaz et al., 2019; Liu et al., 2020). Humans transport species beyond their native ranges to new locations, where they may establish and integrate into local species assemblages (IPBES, 2023; Meyerson & Mooney, 2007). The number of established introduced species increases globally (Seebens et al., 2017, 2021). About ten percent of the established introduced species recorded to date are classified as invasive, i.e., they spread and negatively affect biodiversity, local ecosystems and nature’s contribution to people (IPBES, 2023). Species introductions change the composition of local species assemblages, can contribute to biotic homogenisation and increase extinction risks of native species (Bellard et al., 2021; Díaz et al., 2019). They also affect assemblages of interacting species, such as interacting plants and frugivores, which globally, increasingly share similar species and pairwise interactions (Fricke & Svenning, 2020).

Trophic interactions between frugivorous animals and fleshy-fruited plants are ubiquitous and ecologically crucial, contributing to the provisioning of fruit resources for frugivores and the structure and diversity of plant communities through seed dispersal (Nogales et al., 2024; Rogers et al., 2021). Species morphological traits influence plant-frugivore interactions and seed dispersal. In interactions between fleshy-fruited plants and avian frugivores, the bird species’ bill dimensions correlate with the fruit sizes they handle and swallow, their wing shape influences their flight ability and preferred foraging height, and their body mass determines their energy demand and preferred fruit crop size, a phenomenon termed morphological trait matching (Bender et al., 2018; Dehling et al., 2014; McFadden et al., 2022). Furthermore, large birds might disperse seeds over longer distances than small bird species, due to longer gut retention times and higher flight speeds (Sorensen et al., 2020). Similarly, plant traits, such as height, fruit and seed size, not only determine potential interaction partners of the plant species but also influence fruit removal, gut retention time, and seedling survival (Bracho-Estévanez et al., 2024; Muñoz et al., 2016). The diversity and dominant values of such functional traits in species assemblages influence ecosystem functioning (Gagic et al., 2015; Tilman, 2001).

Global change can alter trait diversity and dominant trait values of species assemblages (e.g., Nowak et al., 2019; Stewart et al., 2022). Yet, our understanding of the consequences of global change for the trait diversity of interacting species is limited (e.g., Nowak et al., 2019). Similarly, the consequences of species’ introductions for trait diversity have mainly been studied within individual taxonomic groups (e.g., Sayol et al., 2021; Tordoni et al., 2019) rather than for assemblages of interacting species. Newly emerging, comprehensive data on biotic interactions and species traits across the globe offer a promising opportunity to address this knowledge gap (e.g., Fricke & Svenning, 2020; Tobias et al., 2022). Here, we leverage these global databases to test how introduced species impact the trait diversity of interacting plant and frugivore species across different locations globally. We address the following research questions:

i. Do introduced species impact trait diversity and dominant trait values within assemblages of interacting plant and frugivore species and what predicts this impact? Introduced species tend to be generalists, e.g., regarding their biotic interactions or habitat preferences (Fristoe et al., 2021; García et al., 2014). Therefore, we expect them to exhibit less extreme trait values than native species, hence decreasing trait diversity. We also predict them to alter dominant trait values in local species assemblages.
ii. Do introduced species affect beta trait diversity across interacting plant and frugivore assemblages? Given that species introductions can cause taxonomic and phylogenetic homogenisation across species assemblages (Daru et al., 2021; Fricke & Svenning, 2020), we anticipate that introduced species will reduce trait differences between assemblages.
iii. Are the trait diversity of bird and plant assemblages related, and do introduced species alter this relationship? Aligning with the observation of morphological trait matching (Dehling et al., 2014; McFadden et al., 2022), we expect a positive relationship of trait diversity and mean bill and fruit size of native bird and plant assemblages. We predict that species introductions disrupt these relationships, given the varying number of introduced species and resulting differences in trait diversity between bird and plant assemblages.

## Material and methods

### Avian frugivore and fleshy-fruited plant assemblages

We based our analyses on a published database providing bipartite networks describing interactions between frugivorous animals and fruit plants globally (Fricke et al., 2022; Fricke & Svenning, 2020). This database reports the introduced status of each species at each location. We selected networks documenting interactions between avian frugivores and fleshy-fruited plants, merging networks recorded at identical latitudes and longitudes to account for temporal or seasonal replication, yielding 216 distinct networks. The species in those networks form the basis for the principal component analyses applied to obtain the two-dimensional PC space within which we estimated trait diversity of species assemblages (see below).

For further analyses, we selected subsets of these 216 networks, including at least two native species (minimum needed to compute native trait diversity) and one introduced species. Networks focusing on preselected bird or plant species subsets were excluded. All species in the dataset are extant. For analyses of the bird assemblages, after removing one outlier introducing bias to the models (Supporting Information), 25 networks remained, including four mainland and 21 island networks, three from the tropics, two from the subtropics, and 20 from higher latitudes (Table S5). For the plant assemblages, 62 networks remained after removing five outliers, including 37 mainland and 25 island networks, 27 from the tropics, 15 from the subtropics, and 20 from higher latitudes (Table S6). The species of one network were considered one assemblage.

### Morphological traits

We compiled measurements of morphological traits influencing interactions between avian frugivores and fleshy-fruited plants via trait matching. For the avian frugivores, we selected bill width and length (mm), which influence the size of fruits bird species can handle and swallow and positively relate to their preferred fruit size (Dehling et al., 2014; Wheelwright, 1985). Additionally, we included wing pointedness measured as hand wing index (i.e., Kipp’s distance divided by wing length multiplied by 100), which influences flight ability and is positively associated with avian foraging height (Dehling et al., 2014; Sheard et al., 2020). Finally, we considered body mass (g), which influences energy requirement and is positively related to the preferred fruit crop mass (Dehling et al., 2014). Bird traits were derived from AVONET (Tobias et al., 2022). Complete trait information was available for all 1271 bird species in the 216 networks.

For the fleshy-fruited plants, we gathered data on fruit length and width (mm), fruit crop mass (g, i.e., mean fresh-fruit mass multiplied by the mean number of ripe fruits of a plant), and plant height (m, a proxy for the height at which fruits are offered). We excluded climbing and epiphytic species from our analyses because their plant height was measured inconsistently and not always represented the height at which fruits are offered. Plant traits were compiled from databases and literature. For some species, trait measurements were gathered from online resources such as electronic floras (Supporting Information). Despite considering multiple data sources, we encountered incompleteness in plant trait data. We compiled data on fruit length for 66%, fruit width for 30%, crop mass for 10%, and plant height for 64% of the 1869 plant species in the 216 networks. Missing plant trait data were phylogenetically imputed (Supporting Information). Since empirical crop mass data was scarce, we considered this trait in the phylogenetic trait imputation but not in the calculations of trait diversity, dominant trait values and beta trait diversity. We calculated mean trait values per species and ln-transformed all traits except hand wing index for further analyses.

### Replication statement

| Scale of inference | Scale at which the factor of interest is applied | Number of replicates at the appropriate scale |
| --- | --- | --- |
| Species assemblages in interaction networks | Species assemblages in interaction networks | 25 assemblages of frugivorous birds, 62 assemblages of fleshy-fruited plants |

### Impacts of species introductions on trait diversity within assemblages

We assessed impacts of introduced species on trait diversity and dominant trait values by calculating differences in trait diversity and mean trait values with and without introduced species for each bird/ plant assemblage. To estimate trait diversity, we followed three steps: First, we ran one principal component (PC) analysis across all 1271 bird species and one across all 1869 plant species in the 216 networks, applying varimax rotation to adjust PC axis orientation. Second, we computed the trait space of each bird and plant assemblage as hypervolume in the resulting two-dimensional PC spaces using Gaussian kernel density and Silverman’s bandwidth estimation (R package hypervolume, Blonder et al., 2014, 2023). Third, we estimated trait diversity as kernel dispersion (method “divergence”), measuring the average distance of a subsample of random points to the centroid of the trait space (R package BAT, Cardoso et al., 2021; Mammola & Cardoso, 2020). This metric reflects the variation of traits in an assemblage and has lower values if species are closer to the centroid and vice versa. We computed kernel dispersion for each assemblage 1000 times and took the average value for further analyses. We calculated trait spaces and kernel dispersion once considering only native species and once including both native and introduced species. To estimate the dominant trait values of the species assemblages, we calculated the mean of each trait per assemblage with and without introduced species.

The impact of introduced species on trait diversity and mean trait values was calculated as the trait diversity/ mean trait value of native and introduced species minus the trait diversity/ mean trait value of only native species in an assemblage. We performed one-sample t-tests across bird/ plant assemblages to assess whether the impact of introduced species on trait diversity and mean trait values differed from zero.

We tested for predictors of the impact of introduced species on trait diversity: a) trait diversity of native species, b) proportion of introduced species within each assemblage (the number of introduced species divided by the total number of species), and c) proportion of introduced species weighted by the level of invasiveness (sum of the level of invasiveness of introduced species divided by the sum of level of invasiveness of all species; details in Supporting Information). For assemblages of fleshy-fruited plants, we additionally included d) percentage of trait coverage (proportion of plant species in the assemblage with empirical trait data).

We fitted linear mixed-effect models to test for relationships between the impact of introduced species on trait diversity and the selected predictors. As some studies sampled several networks at multiple locations, we included the study ID as a random effect in the models. We tested different combinations of predictors and selected the best model based on the Akaike Information Criterion (AIC). For models involving multiple predictors, we assessed multicollinearity with variance inflation factors. Moreover, we tested for spatial autocorrelation in the residuals with Monte-Carlo simulations of Moran’s I (1000 permutations). We performed NULL models to assess whether observed relationships differ from patterns if species were removed randomly from the assemblages (Supporting Information).

### Impacts of species introductions on trait diversity between assemblages

We computed beta trait diversity for each pair of bird and plant assemblages as kernel beta (Cardoso et al., 2021), which quantifies differences between two trait spaces (hypervolumes) based on their overlap and unique parts (Mammola & Cardoso, 2020). We computed beta trait diversity twice based on the hypervolumes computed in the previous step: first, considering only native species and then including native and introduced species. We performed paired t-tests to evaluate differences in beta trait diversity with and without introduced species for birds and plants, respectively.

### Impacts of species introductions on relationships between bird and plant trait diversity

We fitted linear mixed-effect models to test for relationships between bird and plant trait diversity, and mean fruit and bill dimensions, with and without introduced species. Models included percentage of trait coverage as additional predictor and Study ID as a random effect. Spatial autocorrelation in the residuals was tested with Monte-Carlo simulations of Moran’s I. All analyses were performed in R (R Core Team, 2023).

## Results

### Impacts of species introductions on trait diversity within assemblages

Contrary to our expectation (i), the impact of introduced species on trait diversity (measured as the difference with and without introduced species) was not consistently negative. Instead, it varied across avian frugivore assemblages, with some showing decreases and others increases in trait diversity (mean = 0, t = 0.09, df = 24, p = 0.93; Figure 1a). Across plant assemblages, the impact was on average significantly positive (mean = 0.09, t = 3.35, df = 61, p = 0.001, Figure 1b), but also ranged from negative to positive.

**Figure 1:**
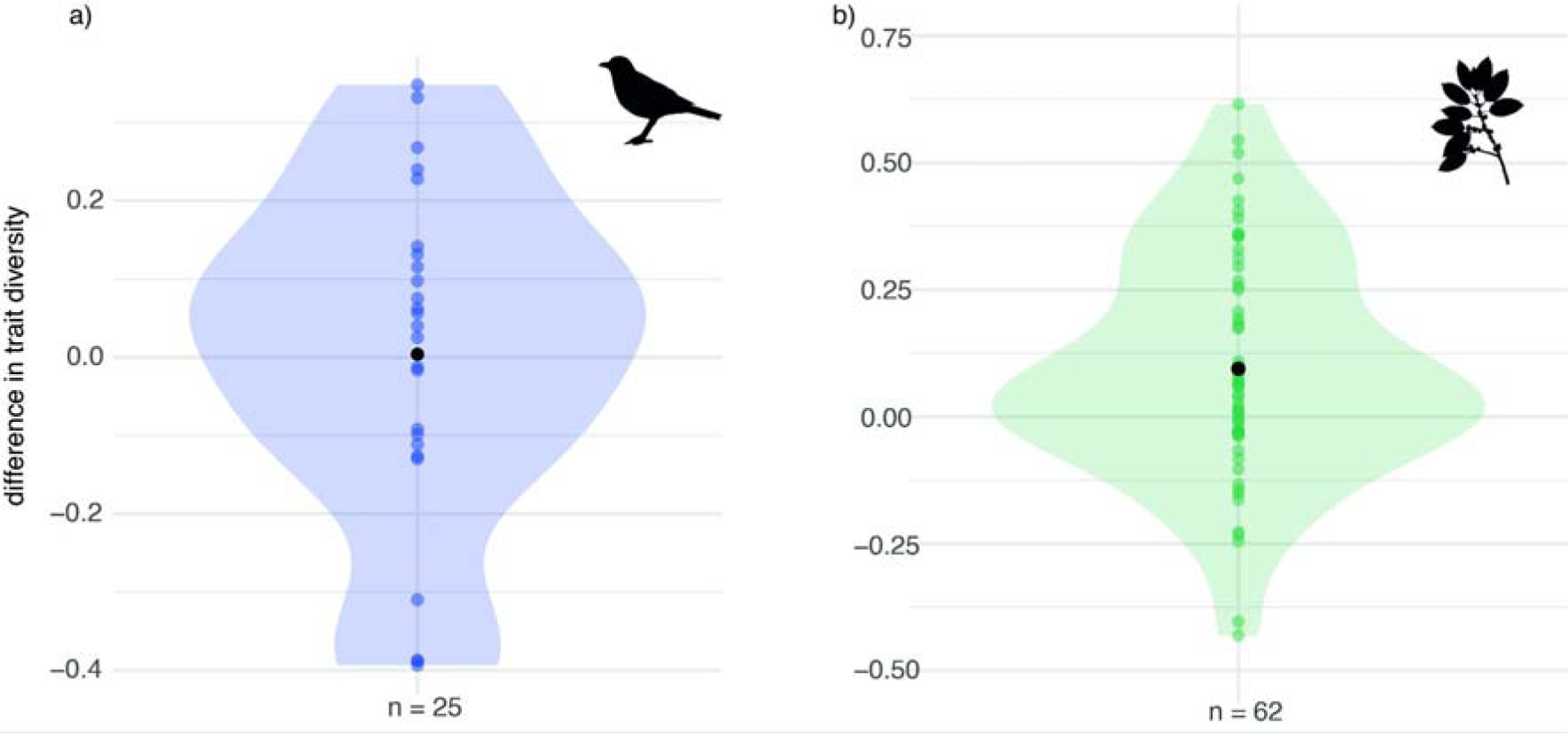
Impact of introduced species on the trait diversity of (a) avian frugivore and (b) fleshy-fruited plant assemblages, measured as the difference in trait diversity with and without introduced species. Coloured points represent individual assemblages, black points denote mean values across assemblages. Coloured areas show the density of the difference in trait diversity. n indicates the number of assemblages analysed.

We examined factors explaining variation in the differences in trait diversity across species assemblages. For avian frugivores, the impact of introduced species on trait diversity was significantly negatively related to trait diversity of native species (estimate = -0.41, se = 0.08, df = 23.87, t = -5.15, p < 0.001, Figure 2a, Table S2a), i.e., introduced species increased trait diversity in assemblages with low native trait diversity but reduced it in those with high native trait diversity. Moreover, the impact of introduced species on avian trait diversity was marginally positively related to the percentage of introduced species weighted by their level of invasiveness (estimate = 0.29, se = 0.15, df = 23.56, t = 1.93, p = 0.07, Figure 2b, Table S2a).

**Figure 2:**
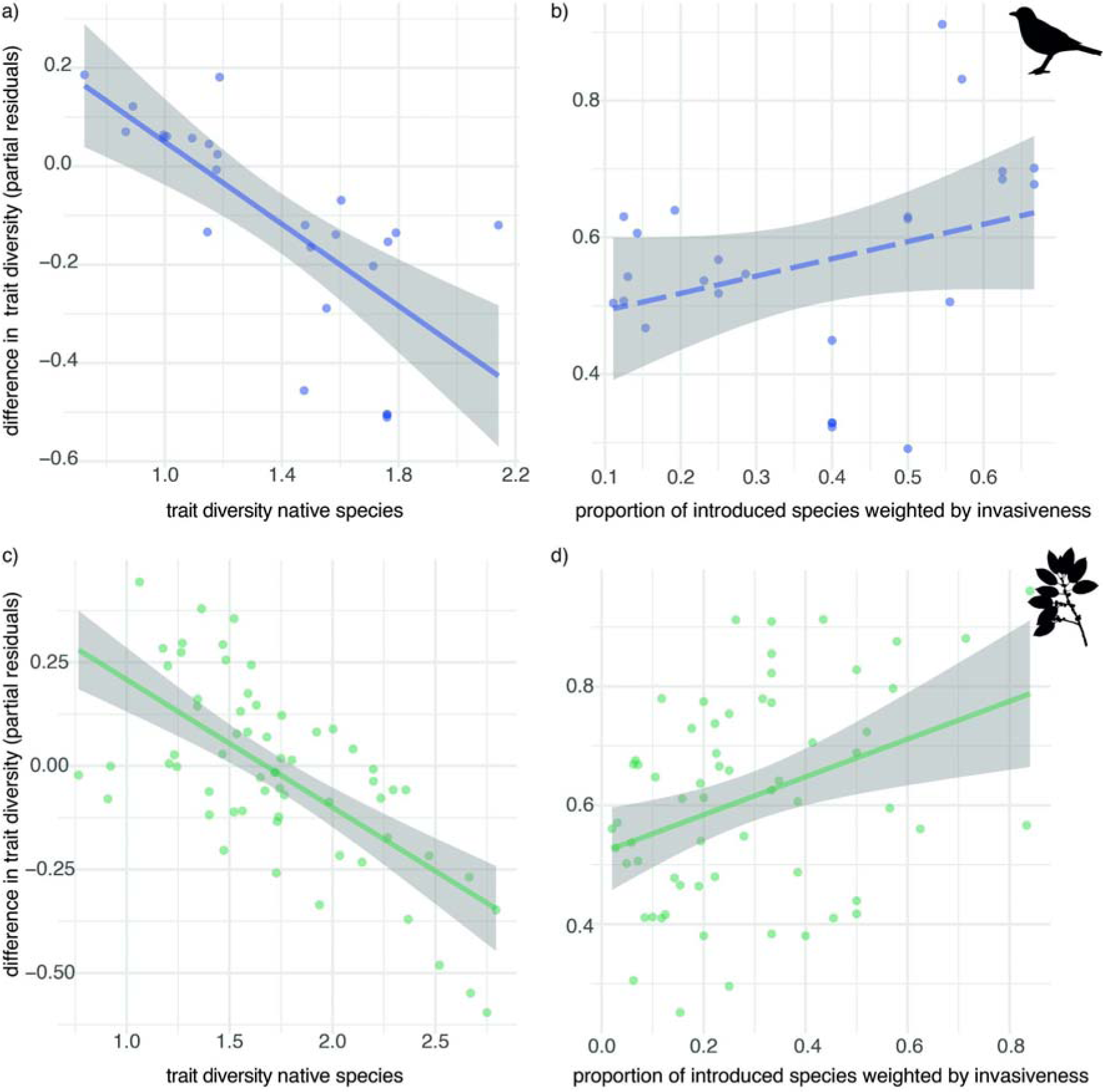
Predictors of the impact of introduced species on the trait diversity of bird (a, b) and plant assemblages (c, d) derived from linear mixed-effect models. Y-axes show partial residuals of the difference in trait diversity. Coloured points represent individual bird/ plant assemblages, lines represent predicted relationships (stippled if 0.05 < p < 0.1), and grey areas depict standard errors.

For fleshy-fruited plants, similar patterns emerged: the impact of introduced species on trait diversity showed a significant negative relationship with native species’ trait diversity (estimate = -0.31, se = 0.05, df = 46.17, t = -6.24, p < 0.001, Figure 2c, Table S2b) and a significant positive relationship with the proportion of introduced species, weighted by their level of invasiveness (estimate = 0.32, se = 0.11, df = 51.61, t = 2.79, p < 0.01, Figure 2d, Table S2b).

The consistent negative relationship between the impact of introduced species on trait diversity and native species’ trait diversity, found for avian frugivores and fleshy-fruited plants, implies that in assemblages with low native trait diversity, introduced species tend to add unique trait combinations, thereby increasing trait diversity. Conversely, in native assemblages with high trait diversity, introduced species tend to reduce the overall trait diversity by adding trait combinations more closely aligning with central native trait values. Null model analyses confirmed that these relationships deviate significantly from scenarios in which species are removed randomly from assemblages (Table S3&4). Additionally, the higher the percentage of introduced species, especially invasive ones, the more likely unique trait values are added, increasing trait diversity.

Across avian frugivore assemblages, we further found a significant increase in mean bill width (difference in mean bill width, mean = 0.12, t = 6.91, df = 24, p < 0.001, Figure 3a), bill length (mean = 0.09, t = 6.65, df = 24, p < 0.001), wing pointedness (mean = 1.69, t = 5.86, df = 24, p < 0.001) and body mass (mean = 0.22, t = 4.52, df = 24, p < 0.00) when comparing bird assemblages with and without introduced species (n = 25).

**Figure 3:**
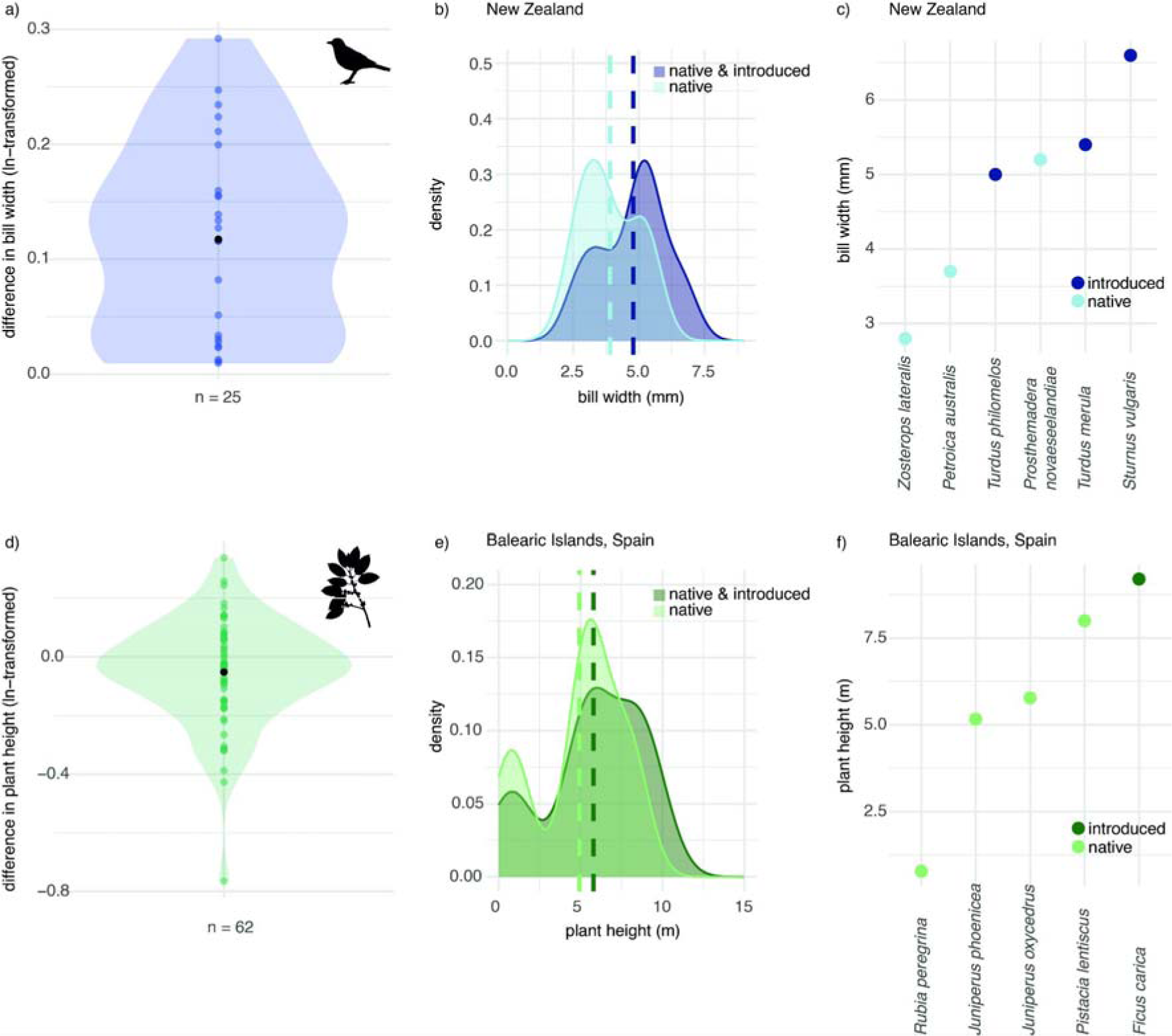
Comparison of individual traits in assemblages with and without introduced species: differences in mean bill width (a) and mean plant height (d) across 25 bird and 62 plant assemblages; differences in bill width (b, c) and plant height (e, f) within one example bird assemblage from New Zealand and one example plant assemblage from the Balearic Islands with full coverage in plant height. Coloured points in (a) and (d) represent species assemblages, black points denote mean values across species assemblages. Density plots in (b) and (e) show differences in trait variation and mean trait values (indicated by stippled lines). Points in (c) and (f) represent individual species and their trait values.

Across the fleshy-fruited plant assemblages, changes in mean trait values were more variable. Mean fruit length significantly increased (mean = 0.06, t = 2.5, df = 61, p = 0.02), changes in mean fruit width were zero (mean = 0.02, t = 1.11, df = 61, p = 0.27) and mean plant height decreased (mean = -0.05, t = -2.28, df = 61, p = 0.03, Figure 3d) across the plant assemblages (n = 62). For all traits, impacts of introduced species ranged from negative to positive. Patterns remained similar when excluding outliers (Supporting Information).

### Impacts of species introductions on trait diversity between assemblages

As predicted (ii), when considering only native species, beta trait diversity was higher than when considering native and introduced species. We found this for avian frugivores (mean difference = -0.15, t = 18.06, df = 299, p-value < 0.001, n = 300 assemblage pairs; Figure 4a) and fleshy-fruited plants (mean difference = -0.05, t = -21.5, df = 1890, p-value < 0.001, n = 1891 assemblage pairs; Figure 4b). These results indicate that species introductions led to assemblages being more similar in their trait combinations.

**Figure 4:**
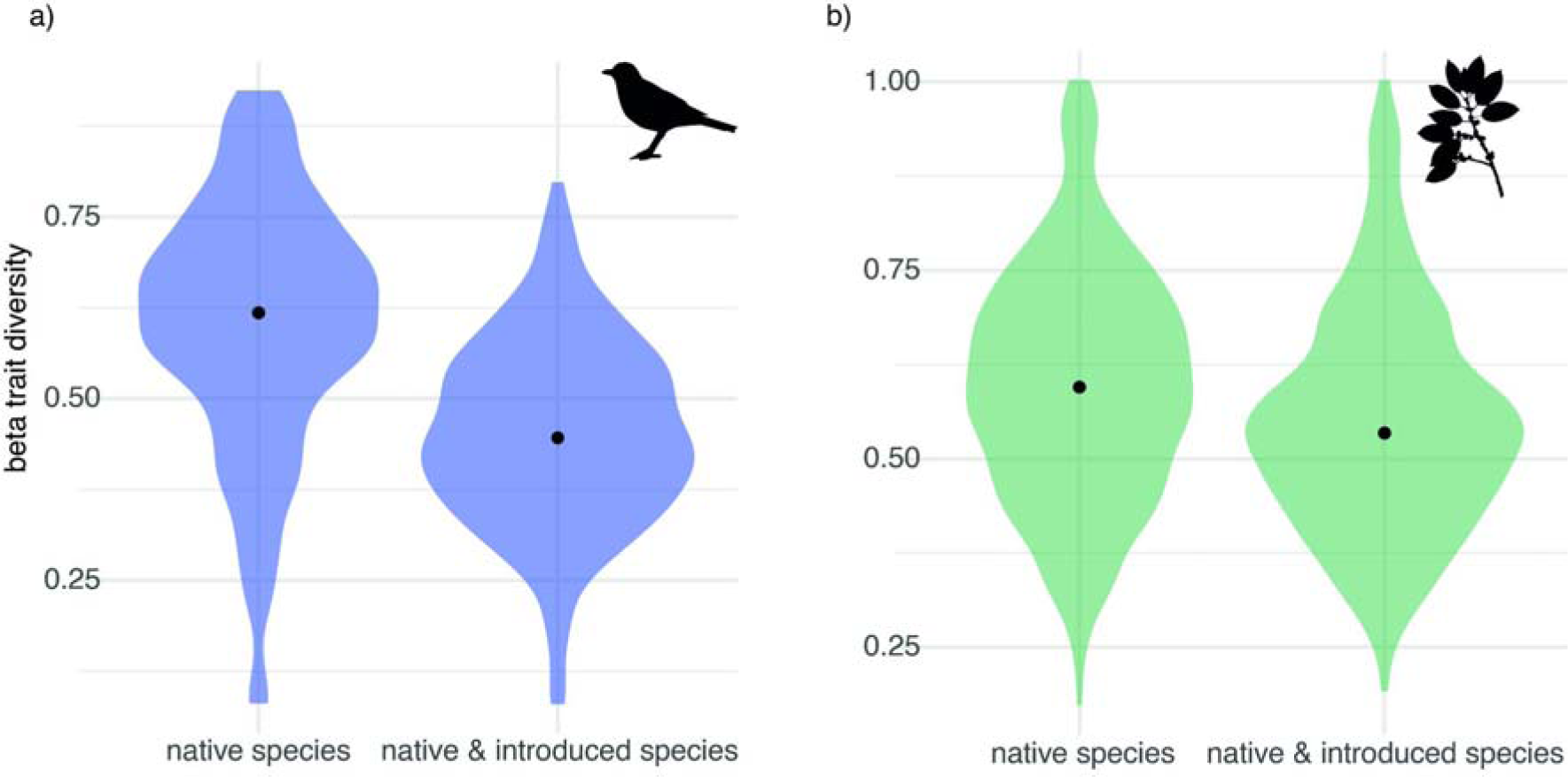
Beta trait diversity comparing assemblages of native species and assemblages of native and introduced species of avian frugivores (a) and fleshy-fruited plants (b). Black points represent mean beta trait diversity across all assemblage pairs. Coloured areas depict density of beta trait diversity.

### Impacts of species introductions on relationships between bird and plant trait diversity

Trait diversity of bird and plant assemblages were not related, neither when focussing on native species (estimate = -0.05, se = 0.14, df = 63.7, t = -0.38, p = 0.70, n = 67 networks with at least one introduced bird or plant species) nor on native and introduced species together (estimate = -0.06, se = 0.14, df = 63.95, t = -0.44, p = 0.66, n = 67). However, mean bill width and fruit width (estimate = 0.48, se = 0.17, df = 44.16, t = 2.9, p = 0.006, n = 67; Moran’s I = -0.07, p = 0.77) and mean bill length and fruit length (estimate = 0.66, se = 0.21, df = 59.94, t = 3.18, p = 0.002, n = 67; Moran’s I = -0.12, p = 0.92) were significantly positively associated when considering native species. Contrary to our prediction (iii), these relationships remained significant when considering both native and introduced species (bill width and fruit width: estimate = 0.57, se = 0.18, df = 51.52, t = 3.21, p = 0.002, n = 67; Moran’s I = -0.06, p = 0.72; bill length and fruit length: estimate = 0.7, se = 0.21, df = 53.03, t = 3.29, p = 0.002, n = 67; Moran’s I = -0.07, p = 0.74). When excluding outliers, results remained qualitatively similar (Supporting Information).

## Discussion

We combined interaction and morphological trait data to study impacts of species introductions on trait diversity of interacting avian frugivores and fleshy-fruited plants at different locations globally. Our findings indicate that introduced species influence trait diversity and dominant trait values within bird and plant assemblages and decrease beta trait diversity across assemblages, highlighting the value of trait data in understanding global change impacts on biotic interactions.

### Native trait diversity influences impacts of introduced species on trait diversity

Contrary to our prediction that introduced species would consistently decrease trait diversity in bird and plant assemblages, the magnitude and direction of the impact of introduced species on trait diversity varied with native trait diversity. Previous studies reported inconsistent results. For example, species introductions increased plant trait diversity on the Canary Islands (Hanz et al., 2022) but decreased it across habitats in Czechia (Loiola et al., 2018). Such variation suggests context dependence, which in ecological studies can be apparent, i.e., due to confounding factors or methodological differences, or related to ecological mechanisms (Catford et al., 2022). Our findings highlight trait diversity of native assemblages as key determinant of functional consequences of species introductions.

Native trait diversity and the similarity between native and introduced species’ traits can influence the integration of introduced species into local assemblages (Hanz et al., 2022; Rojas et al., 2019). Darwin’s naturalisation hypothesis predicts that niche overlap between introduced and native species with similar traits impedes introduced species’ integration into local assemblages. Such biotic resistance to species introductions might be particularly prevalent under high native trait diversity (Hanz et al., 2022; Mathakutha et al., 2019). However, in mutualistic interactions, introduced species with traits matching native preferences may integrate more easily (Rojas et al., 2019; Sperry et al., 2021). Understanding such dynamics in detail requires combining trait data with time series of biotic interactions.

Invasive introduced species might differ in certain traits from non-invasive and native species, explaining why in bird and plant assemblages with a larger proportion of introduced species, particularly species with invasion potential, species introductions increased trait diversity. For example, invasive bird species are often multi-brooded and habitat generalists (Shirley & Kark, 2009), while invasiveness of plants relates to higher values in performance-related traits, such as growth rate and size (van Kleunen et al., 2010). The European Starling (*Sturnus vulgaris*), one of the widest spread introduced species globally (IPBES, 2023), has a relatively large bill compared to other species in assemblages where it has been introduced (e.g., Fig. 3 c). Moreover, fleshy-fruited plant species are often introduced as ornamental or food plants, e.g., the Fig (*Ficus carica*) introduced to the Balearic Islands (Fig. 3 f), potentially explaining their distinct trait values compared to native species.

### Functional consequences of species introductions

In bird and plant assemblages with high native trait diversity, introduced species added trait combinations closer to the average values of the assemblage. However, in assemblages with low native trait diversity, introduced species exhibited greater distinctiveness, increasing trait diversity. The diversity hypothesis predicts that higher trait diversity increases ecosystem functioning by enhancing the complementarity of resource use (Song et al., 2014). For example, morphologically distinct avian frugivores interact with more unique plant species (Pigot et al., 2016), hence, higher bird trait diversity increases the probability that morphologically distinct plant species are being dispersed.

Introduced species might functionally compensate for past species extinctions (García et al., 2014), especially when increasing trait diversity. However, evidence on island birds suggests that functional compensation by introduced species is incomplete (Sayol et al., 2021; Soares et al., 2022; Sobral et al., 2016). Even on Oʻahu, Hawaiʻi, where native plants are dispersed solely by introduced dispersers, these only partially replace extinct native dispersers, primarily dispersing seeds from introduced plant species (Vizentin-Bugoni et al., 2019). Similarly, on Mauritius, introduced species rather act as seed predators than replacing seed-dispersal functions of extinct frugivores (Heinen et al., 2023).

Introduced birds shifted dominant trait values towards larger bills, higher body mass and more pointed wings. Contrastingly, a comparison of historic and modern avian frugivore assemblages on Hawai’i showed that modern assemblages had smaller mean bill width and body mass (Case & Tarwater, 2020). Yet, our study did not consider species extinctions, which often disproportionately impact large frugivores, especially on oceanic islands (Donoso et al., 2017; Heinen et al., 2018). In plant assemblages, changes in dominant trait values were more variable, likely reflecting the broader range of environmental factors shaping plant traits.

Shifts in dominant trait values can influence ecosystem functions (Gagic et al., 2015; Sheng et al., 2023). In plant-frugivore interactions, a shift towards larger birds with larger bill sizes might cause dispersal of larger fruits and seeds (Dehling et al., 2014). Additionally, a shift towards increased body size of seed dispersers might increase chances for long-distance seed dispersal, potentially benefiting gene flow, colonization and climate-tracking ability of plant populations (Donoso et al., 2017; Nowak et al., 2022). Yet, larger-bodied frugivores often have smaller population sizes and are more extinction-prone (Dirzo et al., 2014), possibly limiting their functional contribution. Moreover, introduced frugivores often preferably interact with introduced plants (Fricke & Svenning, 2020; Vizentin-Bugoni et al., 2019). Hence, the composition of plant assemblages may determine the extent to which native plants benefit from introduced dispersers, underscoring the context dependence of introduced species’ functional roles in ecosystems.

### Species introductions reduced beta trait diversity

Our findings of lower beta trait diversity when comparing assemblages with and without introduced species align with observations of decreasing beta diversity of ecological communities in the Anthropocene (Baiser et al., 2012; Li et al., 2020). Anthropogenic drivers, such as landscape simplification, urbanisation, or species introductions, can cause species assemblages to become taxonomically, phylogenetically, or functionally more similar (e.g., Daru et al., 2021; Gámez-Virués et al., 2015; Marcacci et al., 2021). Our results, together with previous findings (Dáttilo et al., 2023; Fricke & Svenning, 2020), indicate that species introductions decrease beta diversity of plant-frugivore assemblages globally, making them more similar in species composition, pairwise interactions, and the composition of traits essential to their biotic interactions.

Reduced beta diversity of traits influencing biotic interactions may homogenise functional roles in ecosystems, here, species’ roles as seed dispersers and food sources, potentially weakening ecosystem resilience by synchronising local responses to large-scale environmental changes (Olden et al., 2004). Furthermore, homogenization of frugivore traits may filter plant traits of interacting species. For example, loss of large frugivores can lead to smaller seed sizes in plant assemblages (Galetti et al., 2013). Hence, declining beta diversity of frugivore traits could cascade to reduce beta diversity of fruit and seed traits.

### Species introductions did not disrupt trait matching

The positive relationship between mean fruit and bill size aligns with the well-established phenomenon of trait matching, which has been observed at the level of individual interactions and species assemblages at different geographical scales (Bender et al., 2018; Dehling et al., 2014; McFadden et al., 2022; Nowak et al., 2019). Unexpectedly, species introductions did not impact this relationship in the assemblages studied here, suggesting no disruption of trait matching between extant species. This indicates that species introductions may not always negatively impact ecological communities, but can also have neutral or positive effects (García et al., 2014; Goodenough, 2010). However, trait matching is only one aspect of intact plant-frugivore interactions. Further factors such as species abundances, their preferences and direct and indirect interactions also require studying.

### Trait-based analyses of global change

Despite mining multiple data sources, significant gaps in plant trait data remained. While phylogenetic trait imputation can fill these gaps, collecting and publishing data on traits influencing biotic interactions (e.g., fruit, seed, flower traits, or animal diets), species abundances and interactions over time, is essential for monitoring functional consequences of global change. Improving intraspecific sampling and geographical coverage are also needed to better capture individual trait variation across space and time (Tobias et al., 2022).

Our results underline the potential of trait-based approaches in understanding impacts of species introductions on biotic interactions and ecosystem functions. By examining effects of introduced species on trait diversity of two interacting trophic levels, we show that while homogenisation of traits is widespread, local impacts depend on the trait composition of recipient communities, the proportion of introduced species and their invasion potential. Understanding such complexities is critical for implementing effective conservation measures and enhancing ecosystem functions like seed dispersal under global change.

## Acknowledgements

LN acknowledges a Feodor Lynen-scholarship from the Alexander von Humboldt Foundation. ID is supported by a Marie Curie Postdoctoral Fellowship (HORIZON-TMA-MSCA-101068643). AT is supported by the Spanish Ministry of Science and Innovation (PID2020-114324GB-C21) and by an ERC AdG (Ref. 101054177). Moreover, we are most grateful to Matthias Schleuning for constructive feedback on the initial ideas for this study. Our warmest thanks to Renske Onstein, who supported us with plant trait data she and her team compiled. We also warmly thank Jörg Albrecht for valuable input on analysing trait diversity with incomplete trait data.

## Author contributions

LN and ID conceived the ideas and designed methodology with input from AT and EF; LN analysed the data with input from ID, EF and AT; LN and ID led the writing of the manuscript. All authors contributed critically to the drafts and gave final approval for publication. Our study is based on published data from different locations across the globe which we cite appropriately in the main manuscript. Our author team is small and covers the countries of Spain, Germany, and the USA. We acknowledge that authors from other study locations, especially from the Global South, could have been included and we plan to improve this in future studies.

## Conflict of interest

The authors declare no conflict of interest.

## Data availability

Data on plant-frugivore interactions, species traits and information on species invasiveness utilised in this study are either published or currently being prepared for publication. All metrics calculated at network-level are included in the supporting information and will be uploaded to a public data repository upon acceptance.

## Supporting Information

### Bird traits

We compiled bird trait data (bill length, bill width, hand wing index and body mass) from AVONET (Tobias et al., 2022). AVONET provides species averages of these traits, averaged over raw morphological measurements on live individuals or museum skins. We matched bird species in the networks and AVONET via the BirdTree taxonomy, as this taxonomy was provided in both datasets.

### Plant traits

We standardised the plant names with the Taxonomic Name Resolution Service to ensure taxonomic compatibility among datasets (see data sources in main manuscript). We only included maximum-like values for plant height. We compiled data on vegetative plant height, seed dispersal unit length, and seed dispersal unit width from TRY. For the TRY data, we focussed on standardised trait values (Column: StdValue) and excluded measurements for which the reported error risk was > 4.0 (Kattge et al., 2011; Kattge & et al., 2020). For a few plant species, individual trait measurements were compiled from the following web pages:

- BIISC. (2022). Big Island Invasive Species Committee. https://www.biisc.org/
- Brisbane City Council. (2022). Weed Identification Tool. https://weeds.brisbane.qld.gov.au/weeds
- CABI Digital Library. (2022). CABI Digital Library. https://www.cabidigitallibrary.org/
- eFloras. (2022a). Flora of China. http://www.efloras.org/flora_page.aspx?flora_id=2
- eFloras. (2022b). Flora of North America. http://www.efloras.org/flora_page.aspx?flora_id=1
- EOL. (2022). Encyclopedia of life. https://eol.org/
- Flora of Pakistan. (2022). Pakistan Plant Database. http://legacy.tropicos.org/projectwebportal.aspx?pagename=Home&projectid=32
- Flora Vascular. (2022). Flora Vascular. https://www.floravascular.com
- González, M. L. G. (2022). Flora Vascular de Canarias. http://www.floradecanarias.com/
- Invasive Plant Atlas of the United States. (2022). https://www.invasiveplantatlas.org/
- Kinsey, B. (2022). Wildlife of Hawaii. https://wildlifeofhawaii.com/
- Missouri Botanical Garden. (2022). Missouri Botanical Garden Plant Finder. http://www.missouribotanicalgarden.org/plantfinder/plantfindersearch.aspx
- National Center for Biotechnology Information. (2022). National Library of Medicine. https://www.ncbi.nlm.nih.gov/
- NParks. (2022). Flora & Fauna Web. https://www.nparks.gov.sg/florafaunaweb/who-we-are
- NSW Government Department of Primary Industries. (2022). NSW WeedWise. https://weeds.dpi.nsw.gov.au/
- Porta. (2022). Porta4U. https://prota.prota4u.org/
- Sistema Nacional de Vigilancia y Monitoreo de plagas. (2022). Base de Datos Fitosanitarios. https://www.sinavimo.gob.ar/
- University of Florida Center of Aquatic and Invasive Plants. (2022). Plant Directory. https://plants.ifas.ufl.edu/plant-directory/
- Useful Tropical Plants. (2022). Useful Tropical Plants Database. https://tropical.theferns.info/
- van Welzen, P. C. (2022). Euphorbiaceae of Malesia. https://www.nationaalherbarium.nl/Euphorbs/

### Phylogenetic trait imputation

To fill gaps in the plant trait data, we performed a phylogenetic imputation of missing traits. To this end, we derived a phylogenetic tree for the 1868 plant species considered in our analysis using the function *phylo.maker* in the R package V.PhyloMaker2 with default settings (Jin & Qian, 2019). We then imputed missing trait values with the function *phylopars* in the R package Rphylopars (Goolsby et al., 2017), selecting a Brownian Motion model of trait evolution. This function reconstructs missing trait values from other observed values of the focal trait weighted by phylogenetic proximity, additionally considering the observed value of other traits through an estimate of the phylogenetic correlation between the traits based on all available observations (Bruggeman et al., 2009).

### Species invasiveness

We derived information on whether an introduced species could be considered invasive or highly invasive from the Global Invasive Species Database by the Invasive Species Specialist Group (Invasive Species Specialist Group, 2022), which also provides a list of 100 of the World’s Worst Invasive Alien Species. We further consulted the CABI Compendium Invasive Species (CABI, 2022) and, in some instances, added information from publications (Cisneros-Heredia, 2018; CONABIO, 2017a, 2017b; Cooke et al., 2019; Dwijayanti et al., 2021). We categorized species as “not invasive” if they were not listed as invasive anywhere, as “invasive” if they were listed as potentially invasive or invasive at a given location, and as “highly invasive” if they were among the 100 worst invasive alien species. This classification is not location-specific but approximates species’ general potential to be invasive. Based on this information, we classified each bird and plant species into one of three levels: native or introduced but not invasive (level of invasiveness = 1), introduced and invasive (level of invasiveness = 2), or introduced and highly invasive (level of invasiveness = 3).

### Outliers

For the final linear mixed-effect models presented in the main manuscript, we removed one outlier for the bird- and five for the plant-side analyses. For the birds, this outlier was one bird assemblage with a particularly high positive difference in trait diversity (> 0.8, see boxplot below), and we wanted to ensure this data point was not driving our results.

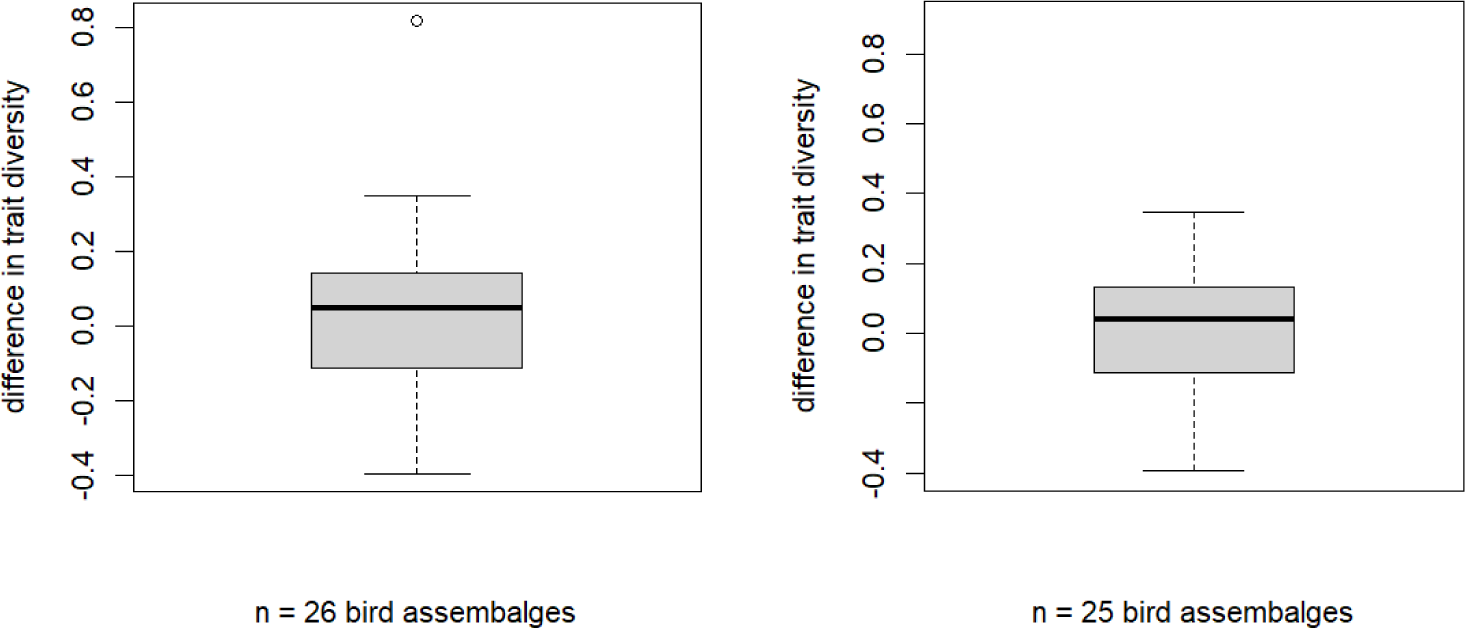

However, when including this outlier in the models, results remained qualitatively similar: the trait diversity of the native bird species was significantly negatively related to the difference in trait diversity (estimate = -0.45, se = 0.07, df = 23.64, t = -6.37, p < 0.001), while the percentage of introduced bird species weighted by their level of invasiveness was significantly positively related to the difference in trait diversity (estimate = 0.38, se = 0.15, df = 24.76, t = 2.62, p = 0.02).

For the plants, outliers were five plant assemblages with a relatively high positive difference in trait diversity (> 0.7 see boxplots below).

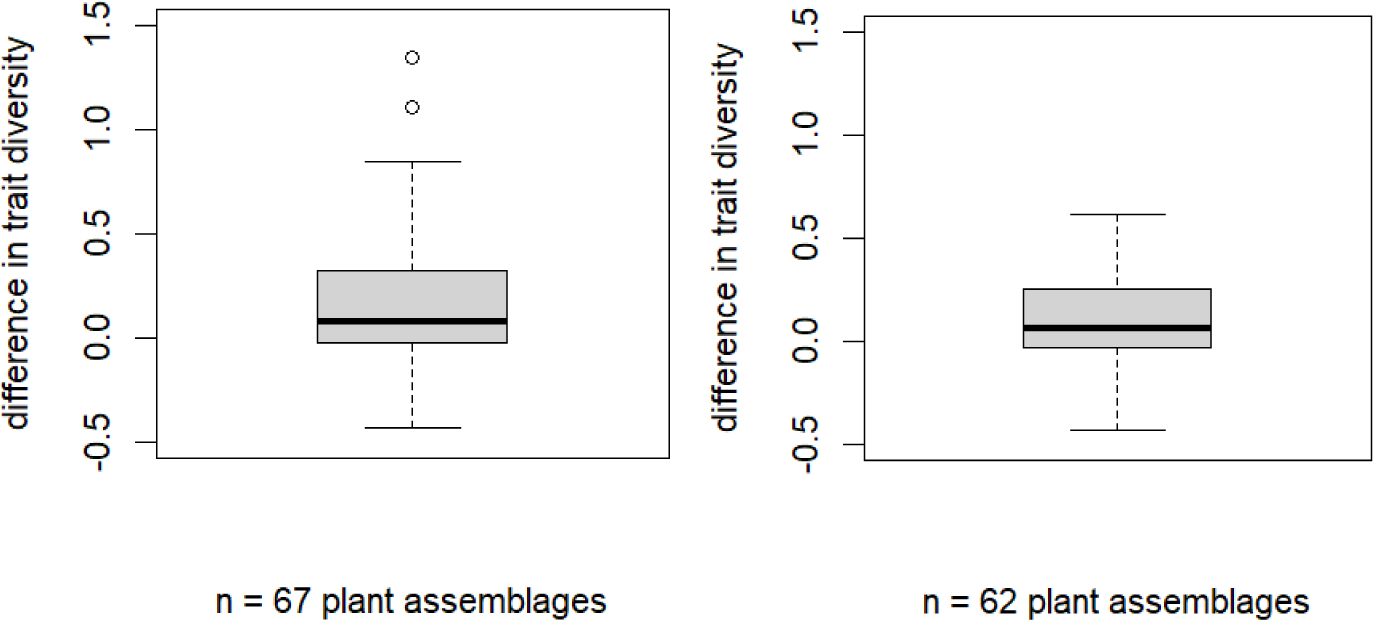

When including these outliers in the model, results remained qualitatively similar: the impact of introduced species on plant trait diversity was significantly negatively related to native trait diversity (estimate = -0.41, se = 0.06, df = 65.43, t = -7.27, p < 0.001, n = 67 species assemblages) and significantly positively to the proportion of introduced species, weighted by the level of invasiveness (estimate = 0.54, se = 0.13, df = 42.04, t = 4.18, p < 0.001, n = 67).

To assess changes in mean trait values of the plant assemblages, we additionally performed t-tests with outliers removed, i.e., one plant assemblage with a change in mean fruit length > 0.8, two with a change in mean fruit width > 0.4, and one with a change in mean plant height of < -0.6. When outliers were removed, patterns remained qualitatively similar: mean fruit length increased (mean = 0.05, t = 2.3, df = 60, p < 0.05, n = 61 plant assemblages), change in mean fruit width did not significantly differ from zero (mean = 0.01, t = 0.3, df = 59, p = 0.8, n = 60), and mean plant height decreased (mean = -0.04, t = -2.02, df = 60, p < 0.05, n = 61).

To test for relationships between mean fruit and bill dimensions, we also fitted mixed-effect models, excluding outliers that might drive model results (four assemblages with a mean bill length >= 3.4 and/or mean fruit length < 1.5; four assemblages with a mean bill width >= 2.1 and/or a mean fruit width < 1). These models revealed similar results to the models presented in the main manuscript: fruit length and bill length were marginally positively related when considering native species only (estimate = 0.44, se = 0.23, df = 57.11, t = 1.9, p = 0.06, n = 63 pairs of interacting bird and plant assemblages) and significantly positively related when considering native and alien species (estimate = 0.63, se = 0.24, df = 53.76, t = 2.69, p < 0.01, n = 63). Fruit width and bill width were significantly positively related for native species only (estimate = 0.34, se = 0.17, df = 46.91, t =2.04, p < 0.05, n = 63) and when considering native and introduced species (estimate = 0.59, se = 0.18, df = 51.18, t = 3.3, p = 0.002, n = 63).

### AIC values of linear mixed-effect models

**Table S1:** AIC values of linear mixed-effect models testing for different combinations of predictors of the difference in trait diversity (Difference FD) of bird and plant assemblages. Predictors tested were: Trait diversity of native species (FD native), proportion percentage of introduced species weighted by their level of invasiveness (% introduced weight.), proportion percentage of introduced species (% introduced), and percentage of plant trait coverage (% trait cov.). The study ID was included as random effect in all models. Details can be found in the main manuscript.

| **Combination of predictors** | **AIC** | **Bird/**<br>**plant** |
| --- | --- | --- |
| **Difference FD ~ FD native + % introduced weight.** | **-**<br>**17.16** | Bird |
| Difference FD ~ FD native + % introduced | -14.71 | Bird |
| Difference FD ~ FD native | -16.22 | Bird |
| Difference FD ~ % introduced weight. | -1.46 | Bird |
| Difference FD ~ % introduced | 0.32 | Bird |
| **Difference FD ~ FD native + % introduced weight. + % trait cov.** | **-**<br>**34.83** | Plant |
| Difference FD ~ FD native + % introduced + % trait cov. | -31.96 | Plant |
| Difference FD ~ FD native + % trait cov. | -29.51 | Plant |
| Difference FD ~ % introduced weight. + % trait cov. | -8.22 | Plant |
| Difference FD ~ % introduced + % trait cov. | -7.61 | Plant |

### Model outputs of best models

**Table S2:** Results of linear mixed-effect models testing for predictors of the impact of introduced species on the trait diversity of avian frugivore (a) and fleshy-fruited plant (b) assemblages. The study ID was included as random effect. The model for birds was based on 25 assemblages, the plant model included 65 assemblages. Reported are the estimates, standard errors (se), degrees of freedom (df), t-values and p-values of both models. Variance Inflation Factors (VIF) indicated no multicollinearity among the predictors, and Moran’s I values (MI) suggested no spatial autocorrelation in the models’ residuals. The models were selected based on their AIC values (Table S1).

|  | estimate | se | df | t-value | p-value | VIF | Moran's I |
| --- | --- | --- | --- | --- | --- | --- | --- |
| a) Birds |  |  |  |  |  |  | MI = 0.11<br>p = 0.11 |
| intercept | 0.47 | 0.12 | 23.89 | 3.79 | 0.001 | na |  |
| Trait diversity native | -0.41 | 0.08 | 23.87 | -5.15 | < 0.001 | 1 |  |
| % introduced species weighted by invasiveness | 0.29 | 0.15 | 23.56 | 1.93 | 0.07 | 1 |  |
| b) Plants |  |  |  |  |  |  | MI = 0.05<br>p = 0.2 |
| intercept | 0.52 | 0.1 | 59.02 | 5.32 | < 0.001 | na |  |
| Trait diversity native | -0.31 | 0.05 | 46.17 | -6.24 | < 0.001 | 1.14 |  |
| % introduced species weighted by invasiveness | 0.32 | 0.11 | 51.61 | 2.79 | < 0.01 | 1.11 |  |
| % trait coverage | 0.06 | 0.08 | 51.21 | 0.68 | 0.51 | 1.06 |  |

### Null models

We performed NULL models to assess whether the relationships between the difference in trait diversity and its predictors differ from patterns we would find, if random species would be removed from the assemblages, i.e., to verify that the patterns we find are indeed due to the species introductions. To this end, we randomly removed the same number of species as introduced species from each analysed bird assemblage and computed trait diversity of the resulting random assemblages. We subtracted these random assemblages’ trait diversity from the complete assemblages’ trait diversity, including all native and introduced species. We then fitted linear mixed-effect models to test for relationships between this difference and trait diversity of native species and the percentage of introduced species weighted by the level of invasiveness. We included the study ID as a random effect in the linear mixed-effect model. We repeated this procedure 10,000 times and extracted the model coefficients for trait diversity and the percentage of introduced species weighted by the level of invasiveness for each repetition. We compared the 2.5 and 97.5 % quantiles of the model coefficients from the 10,000 randomisations to the coefficients of the models testing for predictors of the impact of introduced species on bird trait diversity with and without introduced species. If the latter coefficients lay beyond these quantiles, the patterns were interpreted as non-random.

**Table S3:**
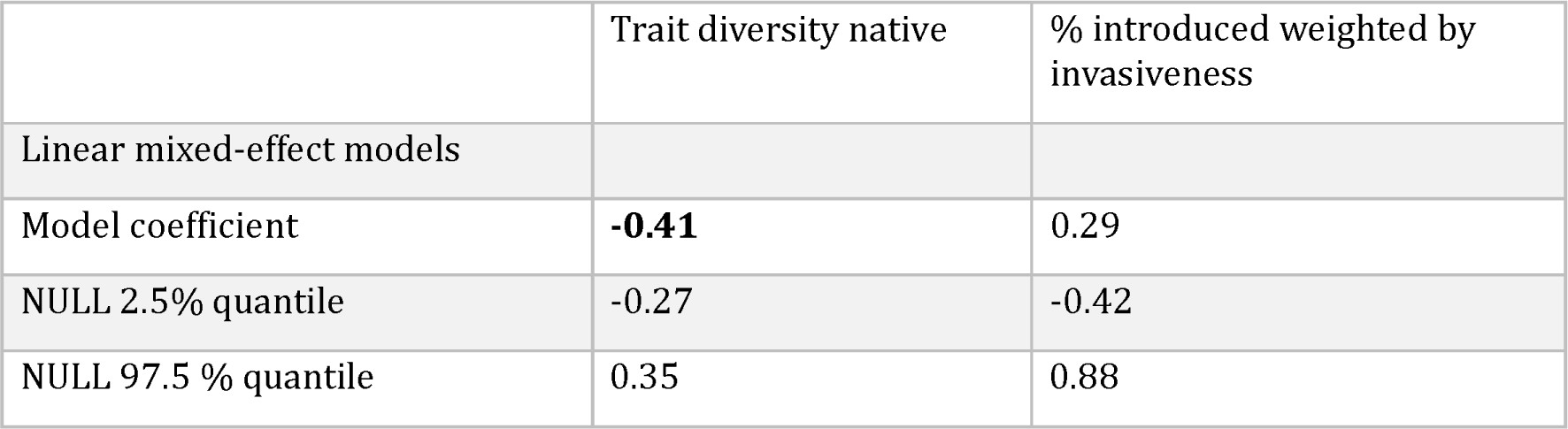
Results of NULL models performed for birds. The model coefficient of the models testing for predictors of the impact of introduced species on bird trait diversity with and without introduced species and the 2.5 and 97.5 % quantiles of the coefficients from the 10,000 runs are reported. Values in bold indicate where the model coefficient is larger/smaller than those quantiles.

**Table S4:** Results of NULL models performed for plants. The model coefficient of the models testing for predictors of the impact of introduced species on bird trait diversity with and without introduced species and the 2.5 and 97.5 % quantiles of the coefficients from the 10,000 runs are reported. Values in bold indicate where the model coefficient is larger/smaller than those quantiles.

|  | Trait diversity native | % introduced weighted by invasiveness |
| --- | --- | --- |
| Linear mixed-effect models |  |  |
| Model coefficient | **-0.31** | 0.32 |
| NULL 2.5% quantile | -0.14 | -0.33 |
| NULL 97.5 % quantile | 0.12 | 0.62 |

### Network-level information

**Table S5:**
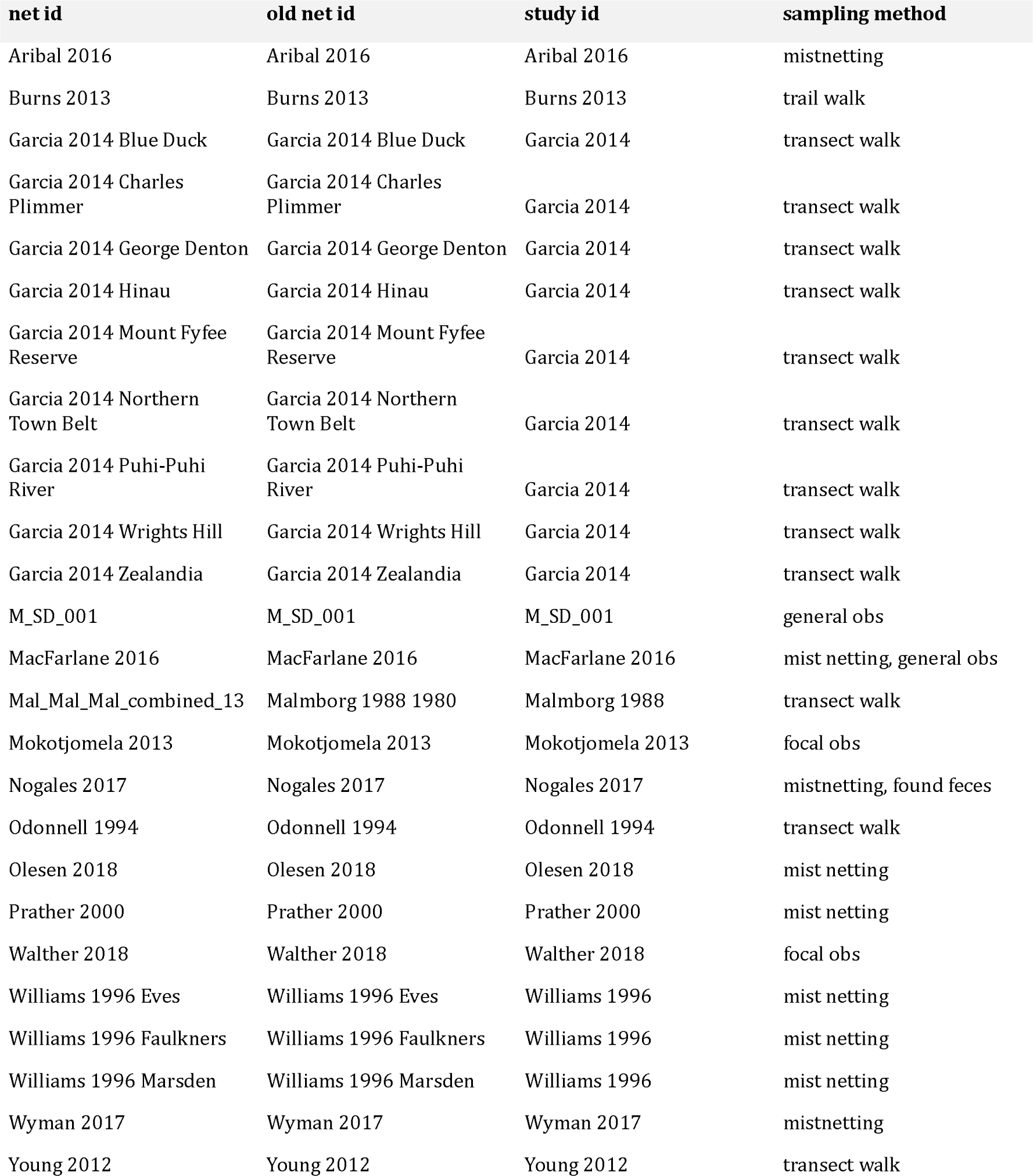
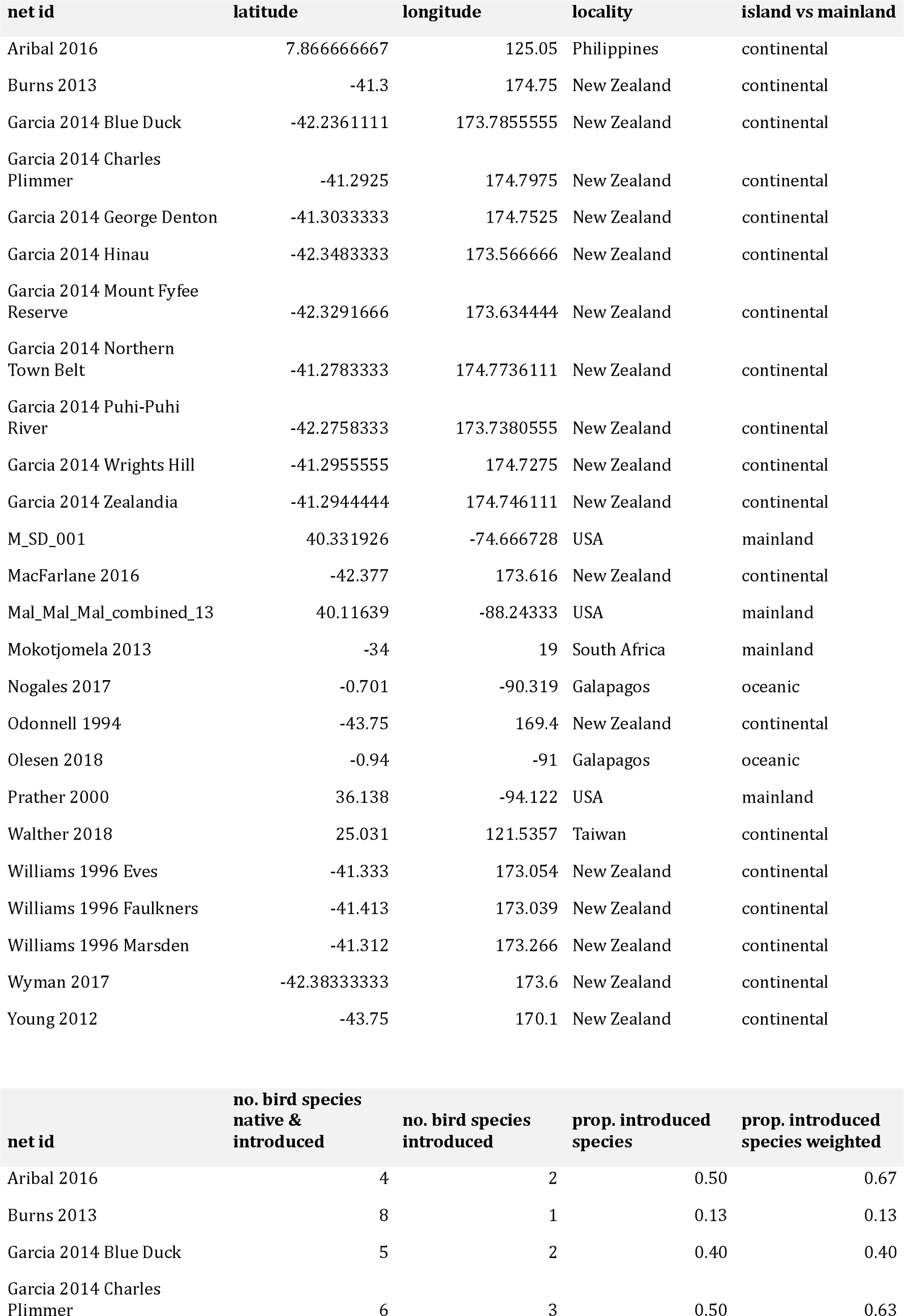
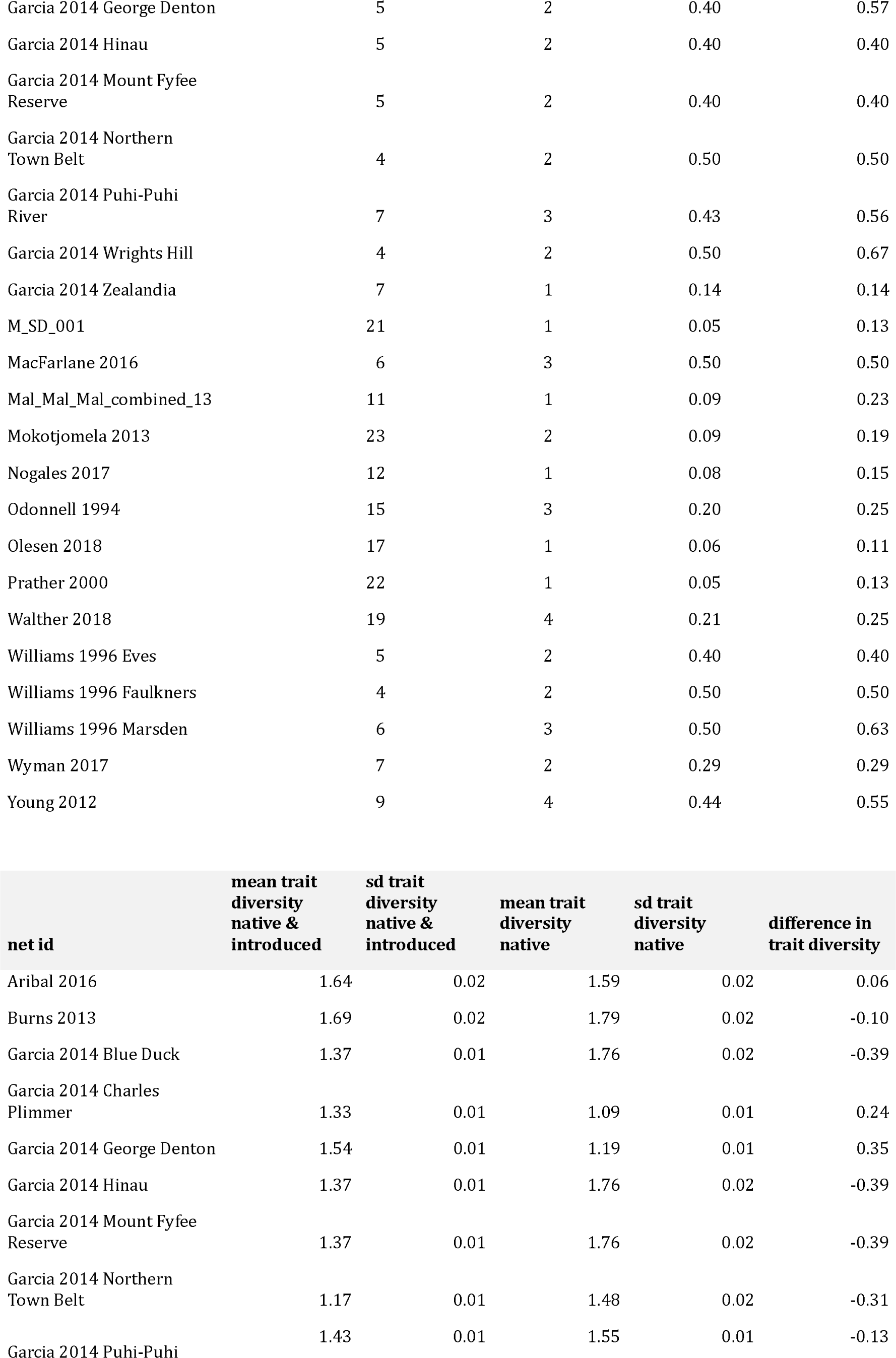
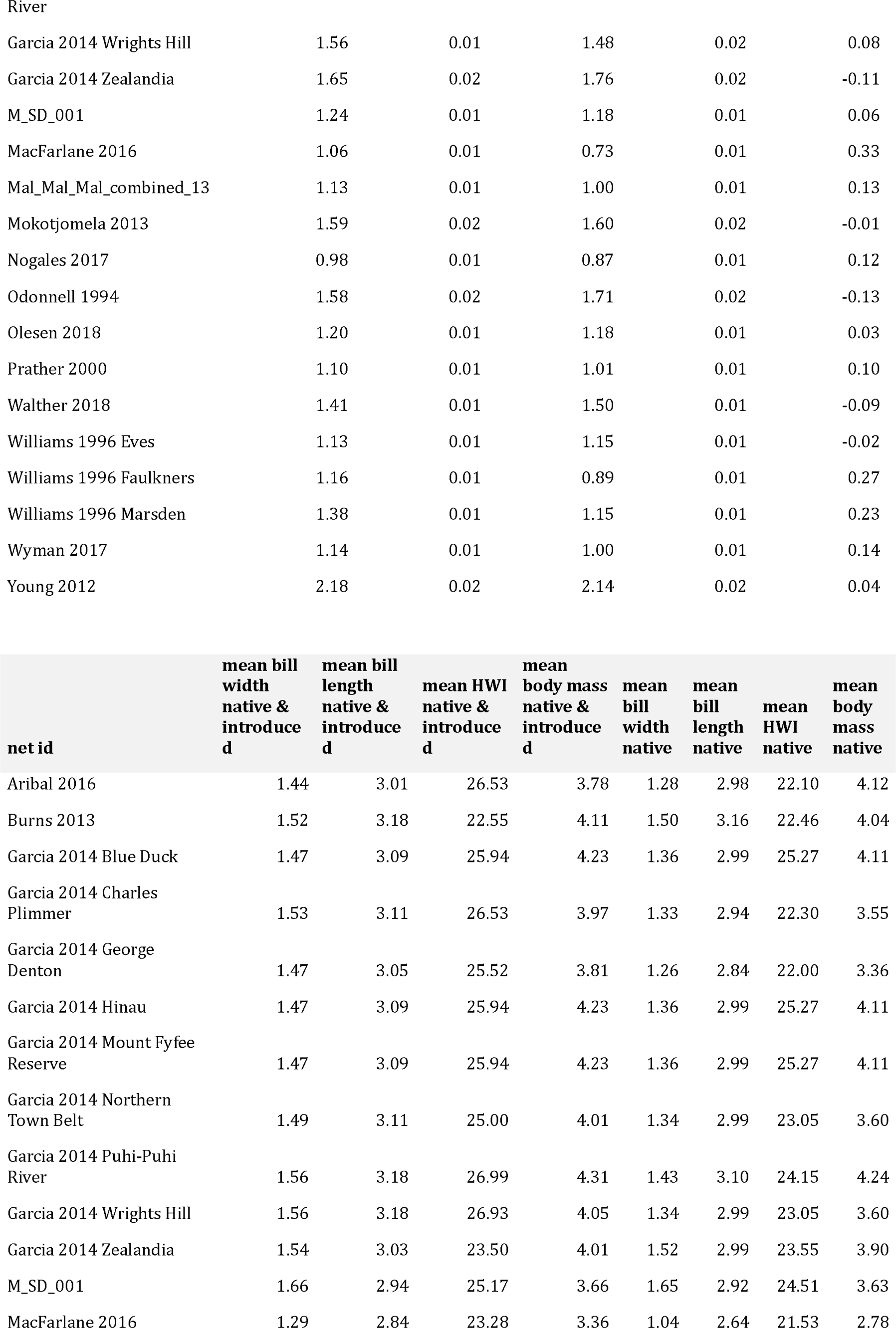
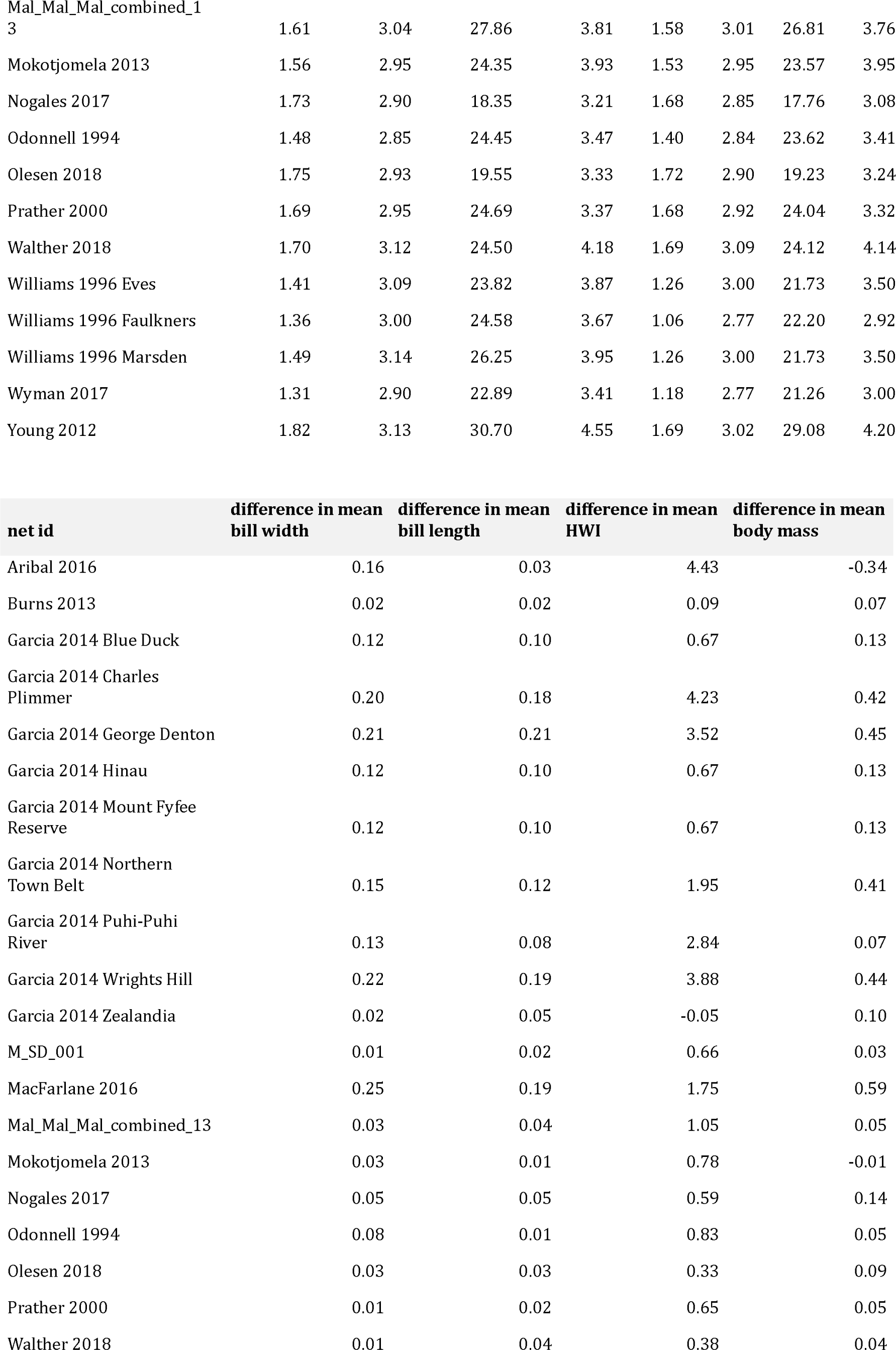
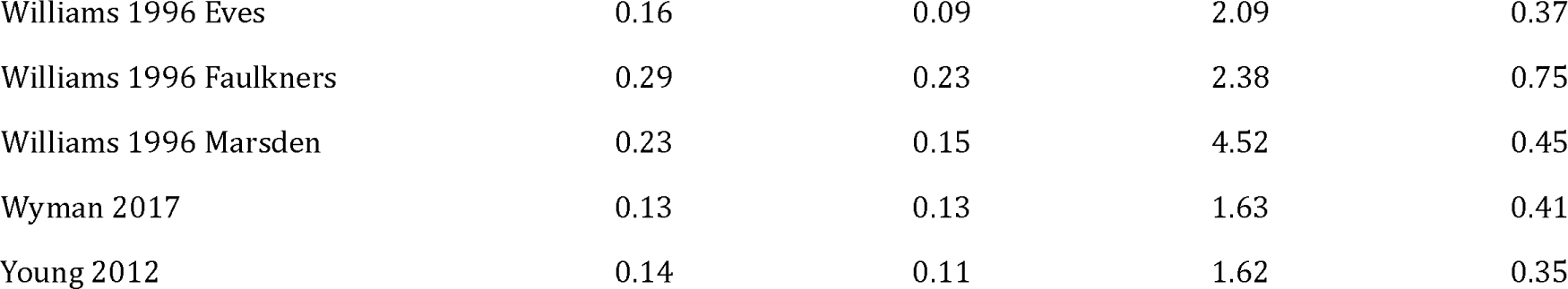
Network-level information and calculations for the networks utilised for the analyses of the frugivorous bird assemblages. Given are the network identifier (*net id*), the old network identifier (*old net id*) before merging temporal replicates of the same network (similar identifiers utilized in the original dataset by (Fricke et al., 2022; Fricke & Svenning, 2020)), the *study id* which indicates the study the networks were collected for (similar identifiers utilized in the original dataset by (Fricke et al., 2022; Fricke & Svenning, 2020)), the *sampling method*, the *latitude* and *longitude* at which networks were recorded, the *locality* of the network, whether the network is located on an oceanic or continental island or mainland (*island vs mainland*), the number of native and introduced bird species (*no. bird species native & introduced*), the number of introduced bird species (*no. bird species introduced*), the percentage of introduced bird species (*prop. introduced bird species*), the percentage of introduced bird species weighted by the level of invasiveness (*prop. introduced bird species weighted*), trait diversity of native and introduced bird species given as mean kernel dispersion across the 1000 runs (*mean kernel dispersion native & introduced*) and the standard deviation of kernel dispersion across the 1000 runs (*sd kernel dispersion native & introduced*), trait diversity of native bird species (*mean kernel dispersion native, sd kernel dispersion native*), the difference in trait diversity with and without introduced species (*difference in kernel dispersion*), the *mean bill width*, *bill length*, *hand wing index* (HWI) and *body mass* of native and introduced species and of native species and the difference in these traits with and without introduced species, all traits except HWI were ln-transformed prior to the analyses.

**Table S6:**
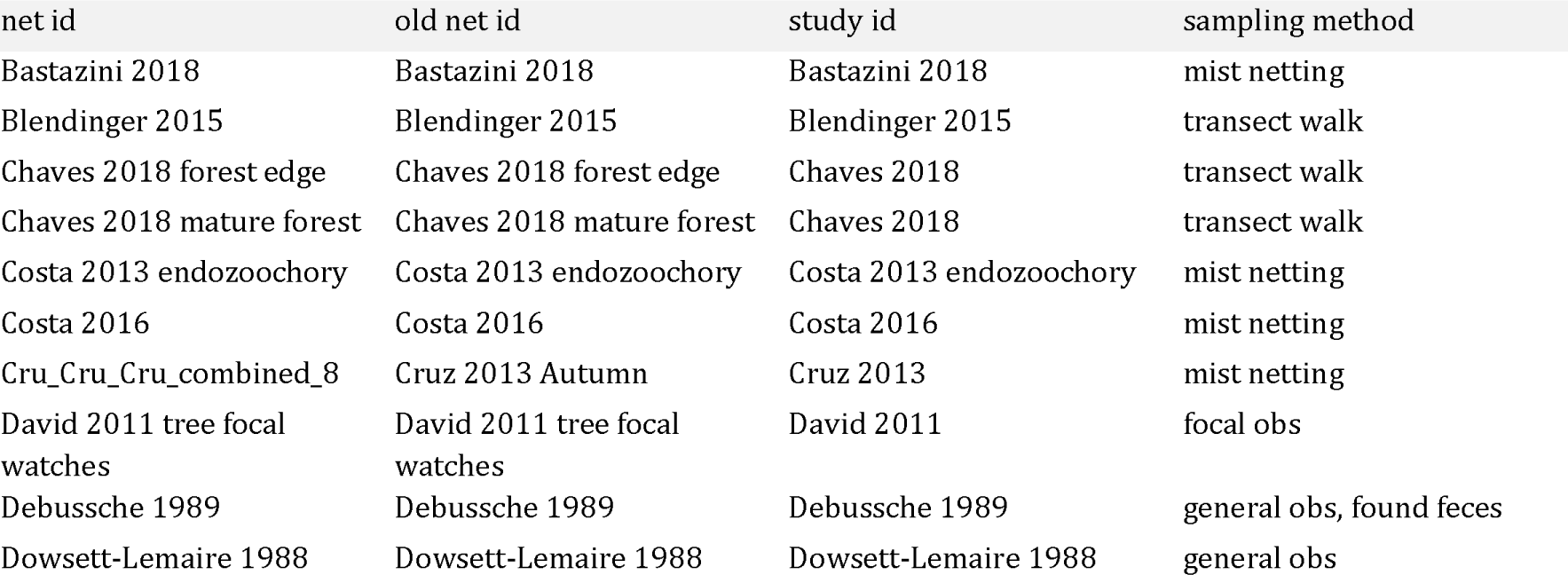
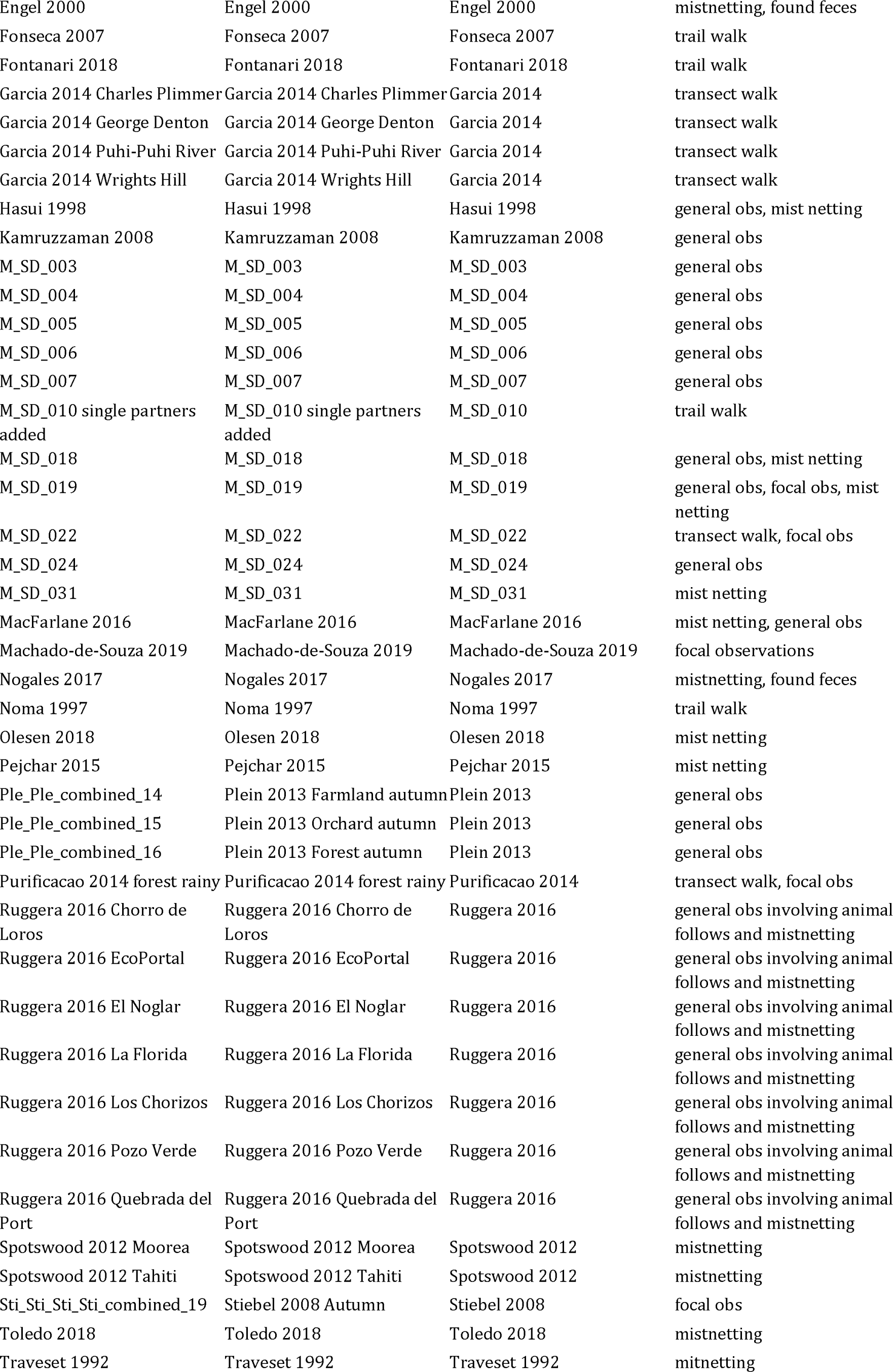
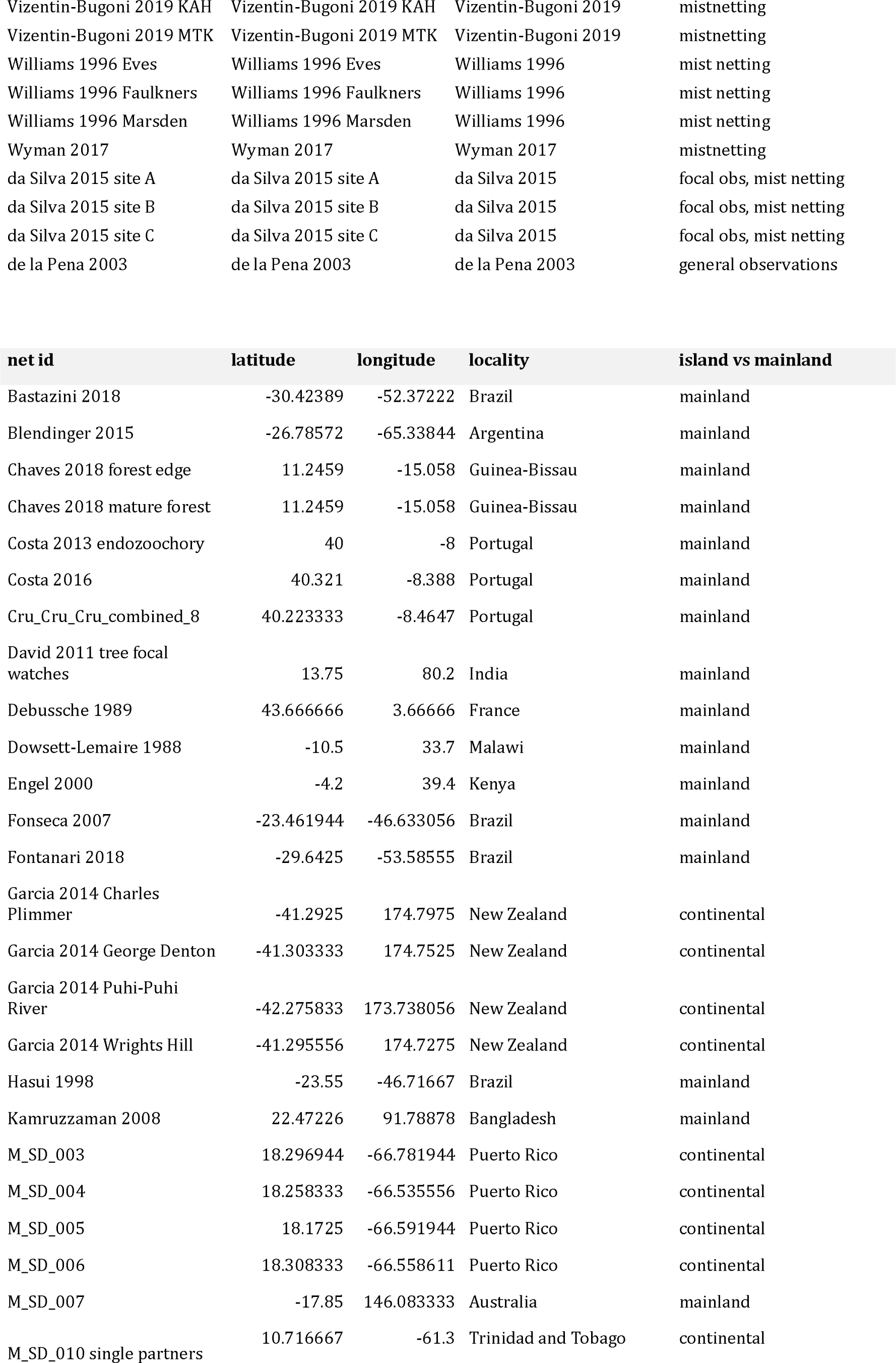
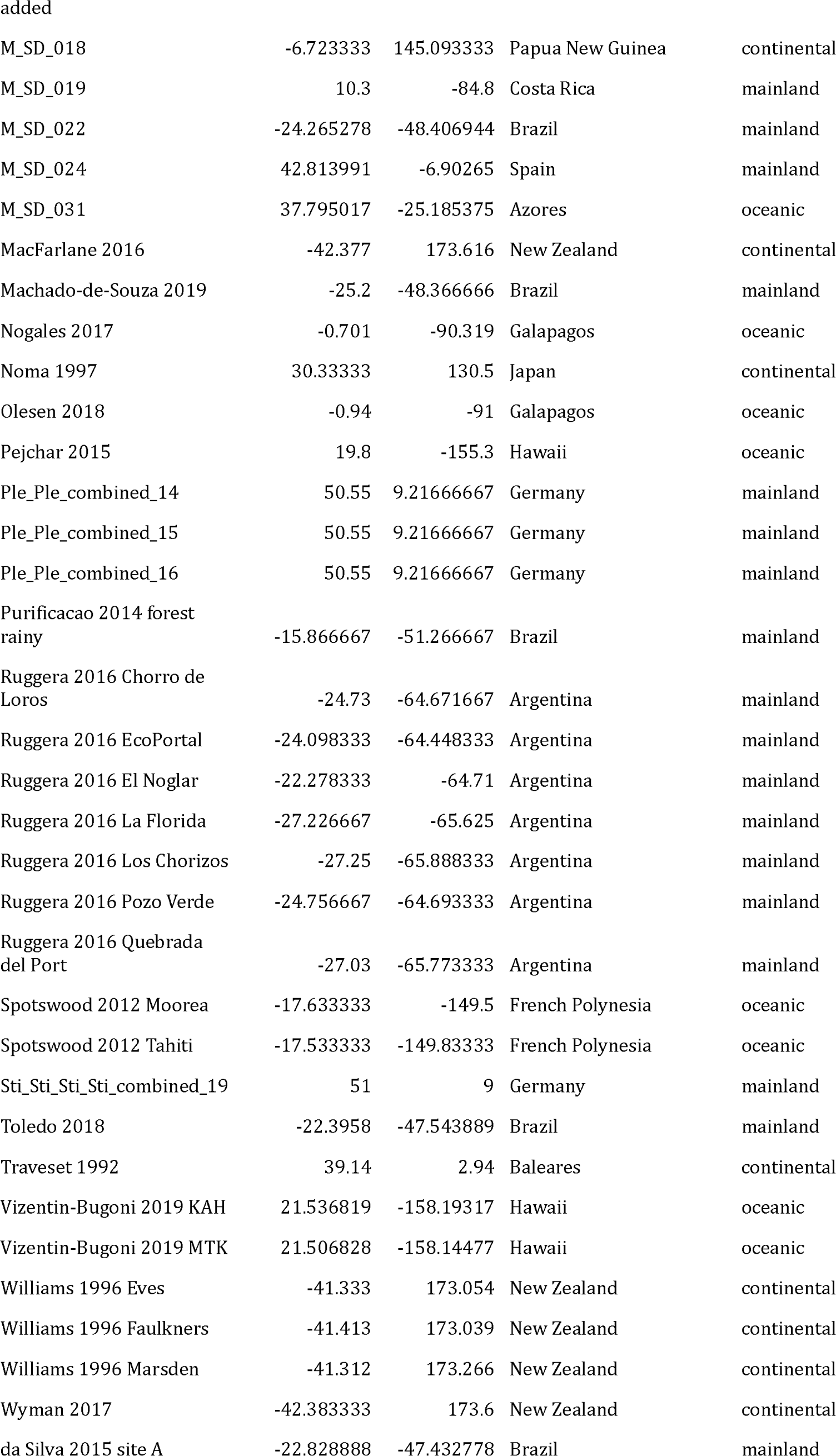
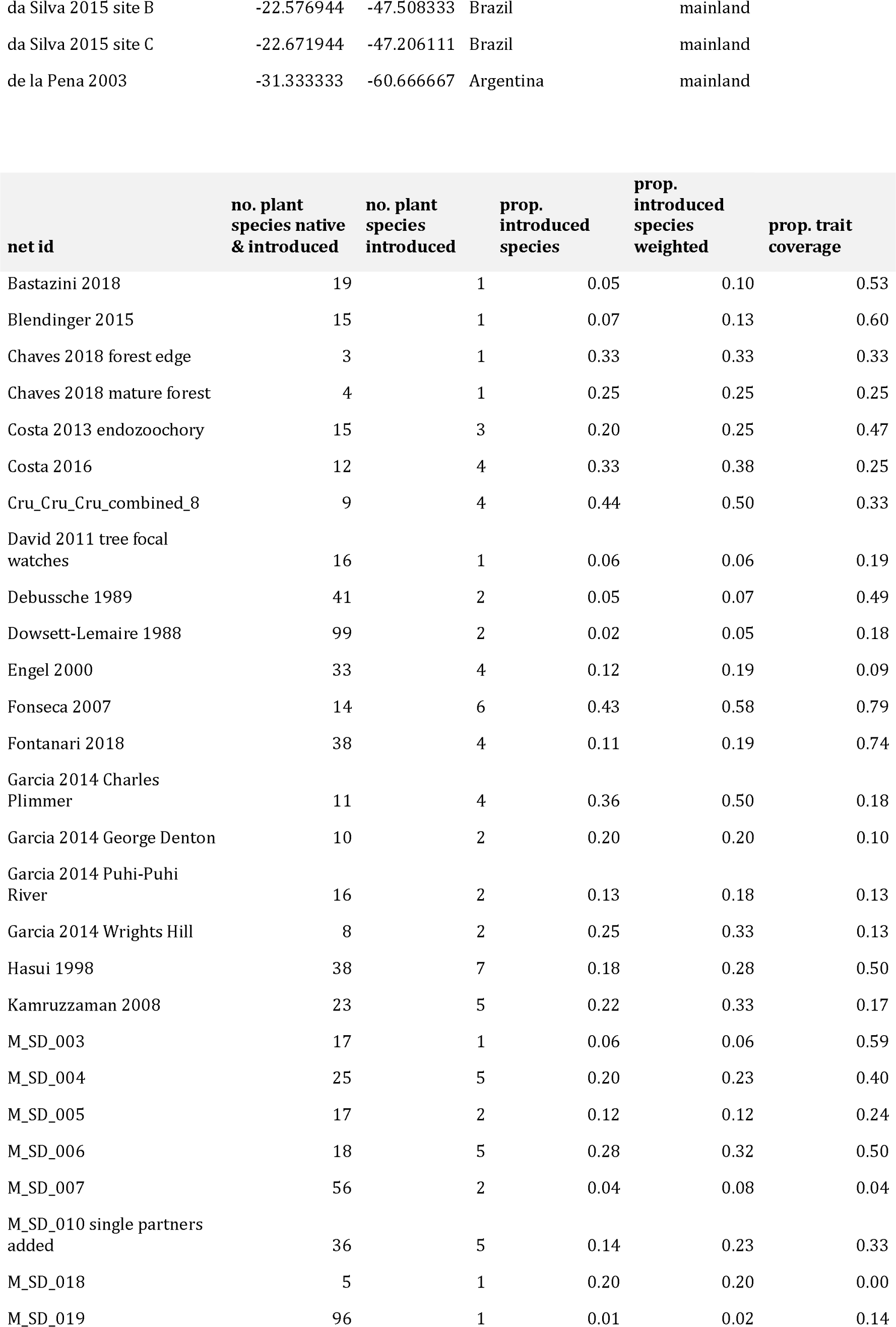
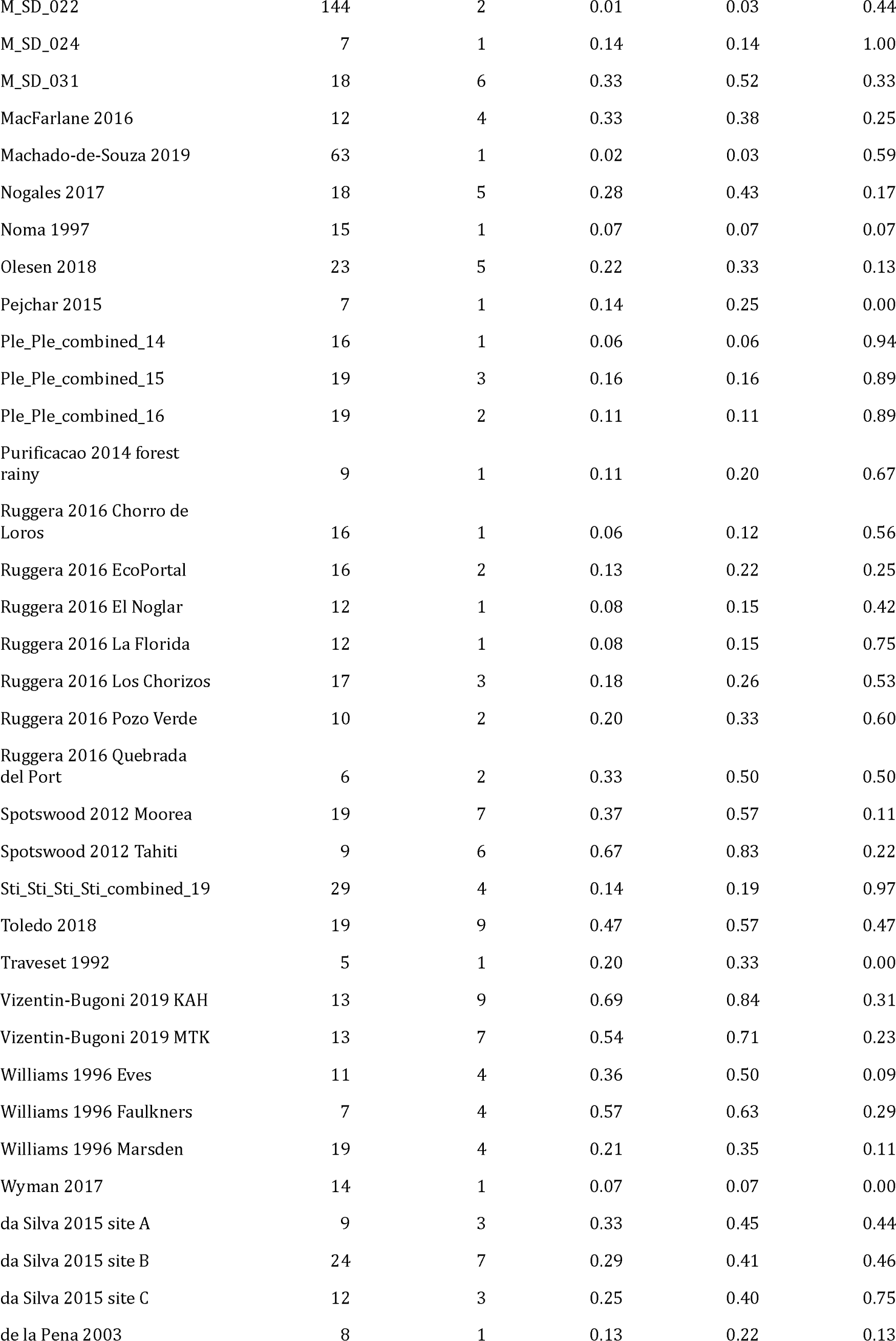
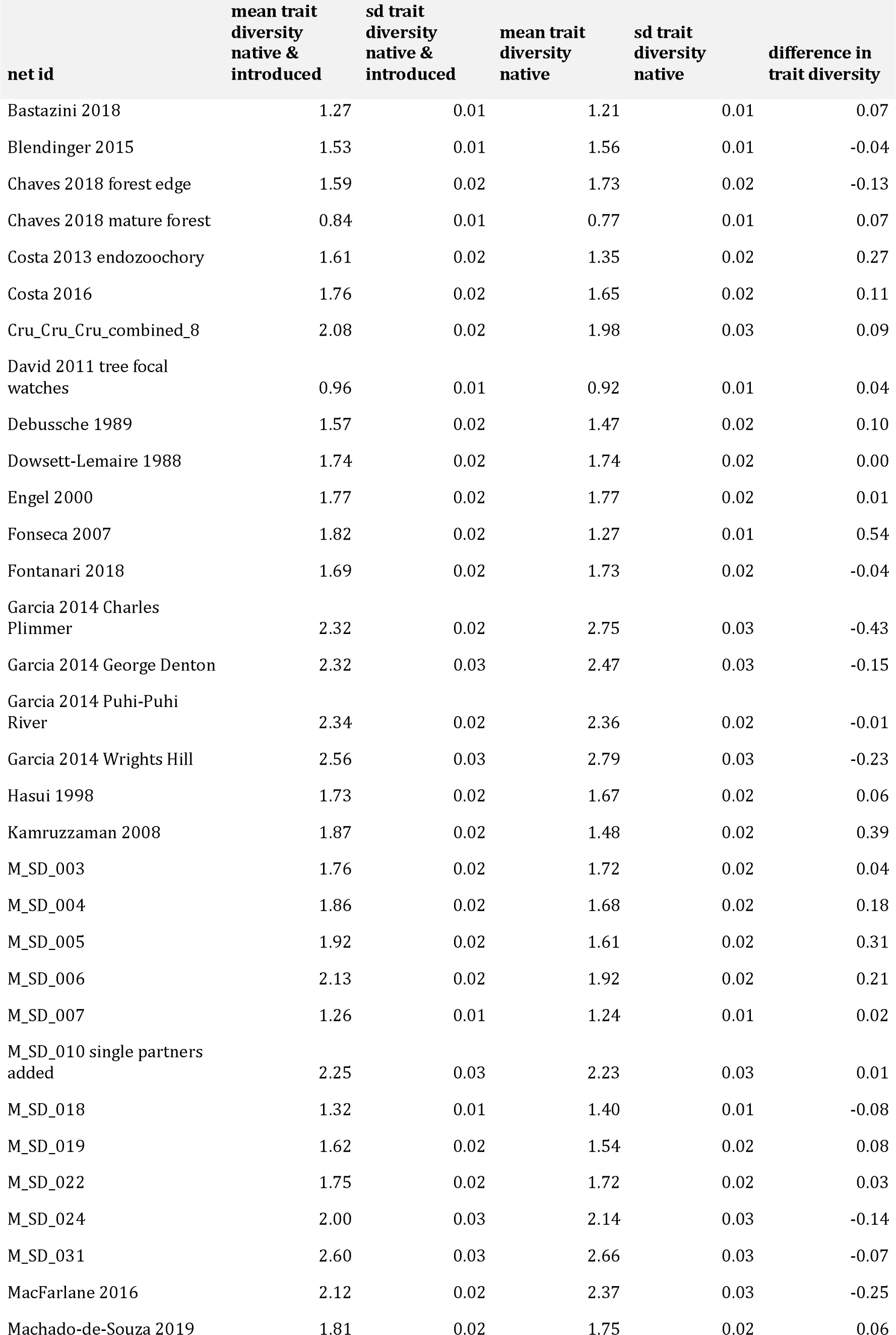
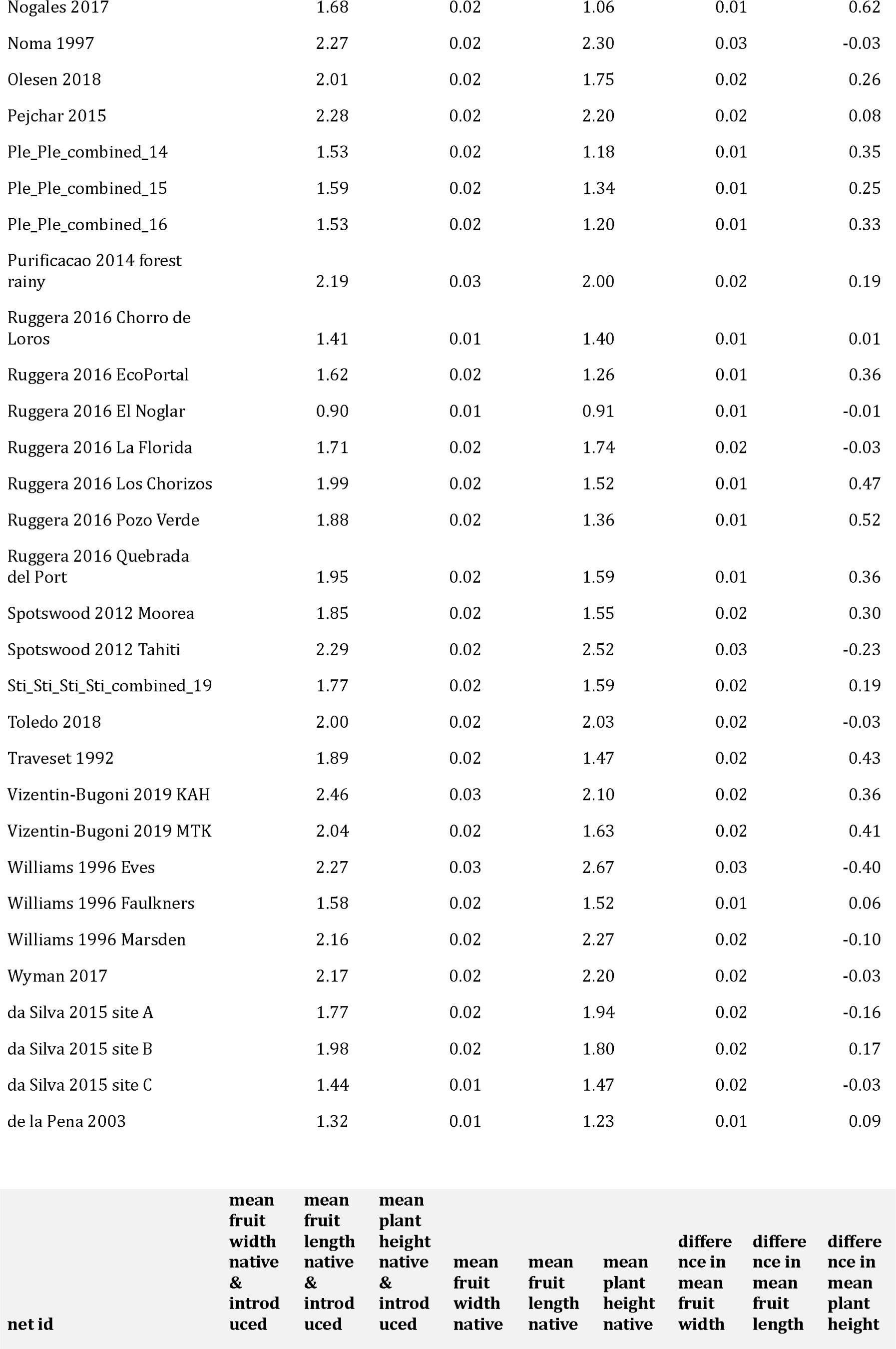
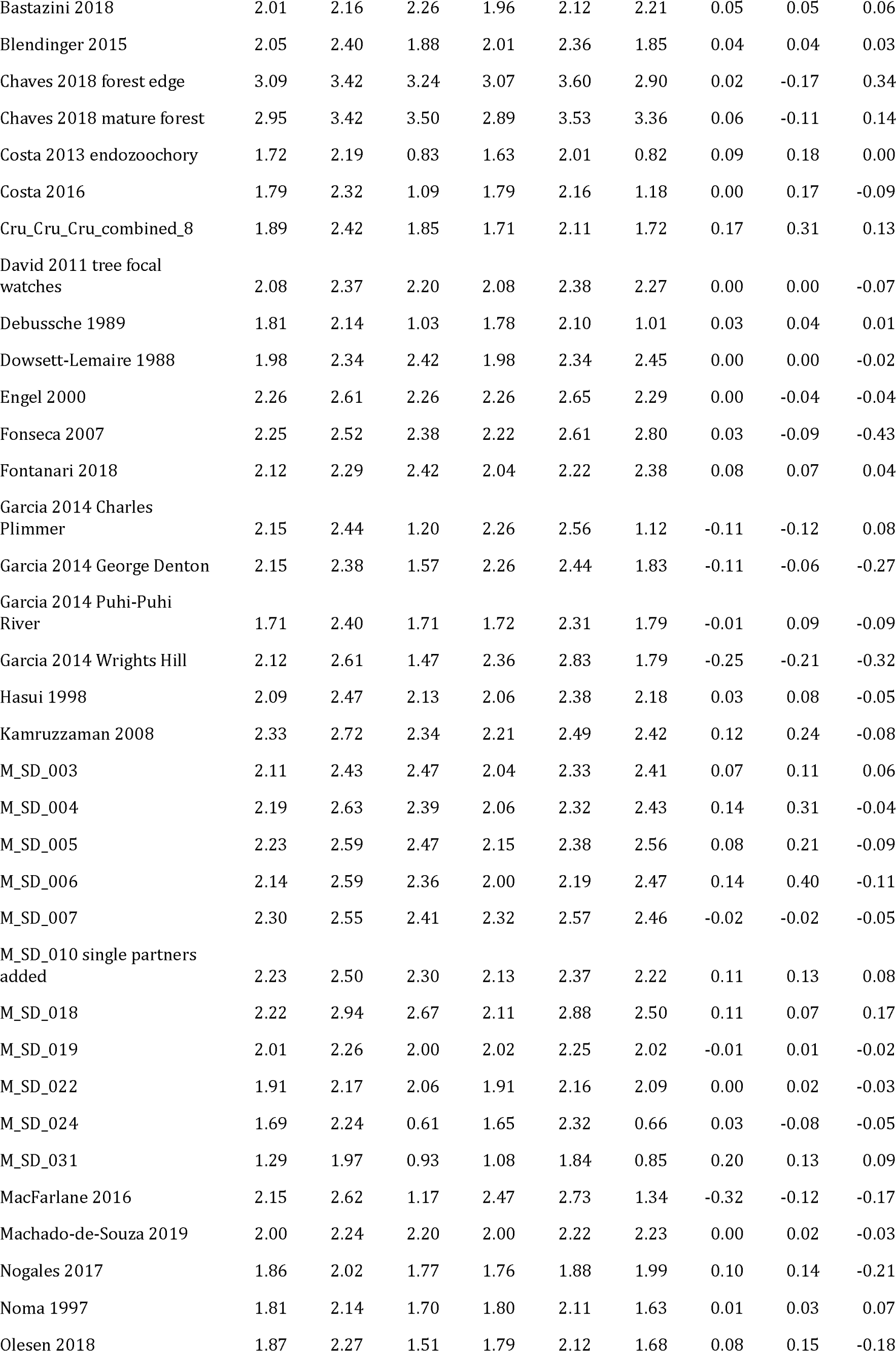
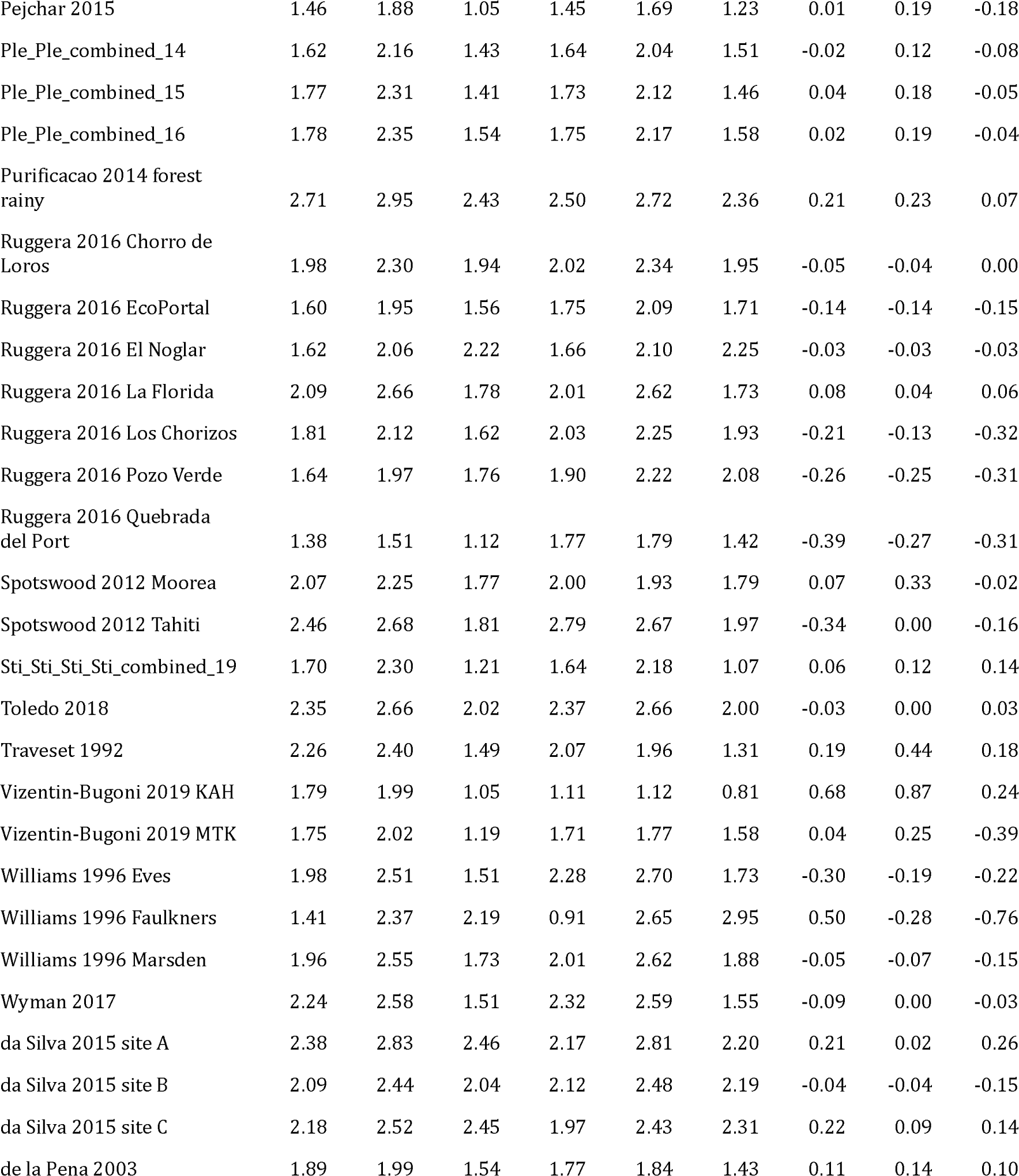
Network-level information and calculations for the networks utilised for the analyses of the fleshy-fruited plant assemblages. Given are the network identifier (*net id*), the old network identifier (*old net id*) before merging temporal replicates of the same network (similar identifiers utilized in the original dataset by (Fricke et al., 2022; Fricke & Svenning, 2020)), the *study id* which indicates the study the networks were collected for (similar identifiers utilized in the original dataset by (Fricke et al., 2022; Fricke & Svenning, 2020)), the *sampling method*, the *latitude* and *longitude* at which networks were recorded, the *locality* of the network, whether the network is located on an oceanic or continental island or the mainland (*island vs mainland*), the number of native and introduced plant species (*no. plant species native & introduced*), the number of introduced plant species (*no. plant species introduced*), the proportion of introduced plant species (*prop. introduced plant species*), the proportion of introduced plant species weighted by the level of invasiveness (*prop. introduced plant species weighted*), the trait coverage (*prop. trait coverage*), the trait diversity of native and introduced plant species given as mean kernel dispersion across the 1000 runs (*mean kernel dispersion native & introduced*) and the standard deviation of kernel dispersion across the 1000 runs (*sd kernel dispersion native & introduced*), the trait diversity of native plant species (*mean kernel dispersion native, sd kernel dispersion native*), the difference in trait diversity with and without introduced species (*difference in kernel dispersion*), the mean *fruit width*, the *mean fruit length*, and the *mean plant height* of native and introduced species, and of native species, and the difference in these traits with and without introduced species (all traits were ln-transformed prior to the analyses).

## Notes

### Competing Interest Statement

The authors have declared no competing interest.

